# The Role of EBP50 in Regulating Endothelial-To-Mesenchymal Transition in Pulmonary Hypertension

**DOI:** 10.1101/2022.05.31.494181

**Authors:** Anastasia Gorelova, Mariah Berman, Maryam Sharifi-Sanjani, Andres Pulgarin Rocha, Anas Alsuraimi, Patricia Riva, Vinny Negi, Thomas Bertero, Dmitry Goncharov, Tatiana Kudryashova, Atinuke Dosunmu-Ogunbi, Arun Rajaratnam, Lloyd Harvey, Mehdi Nouraie, Jeffrey Baust, Timothy Bachman, Andrea Sebastiani, John Sembrat, Sara O. Vargas, Aaron B. Waxman, Rajan Saggar, Rahul Kumar, Brian Graham, Adam C. Straub, Elena A. Goncharova, Ana Mora, Stephen Y. Chan, Imad Al Ghouleh

**Affiliations:** Department of Pharmacology and Chemical Biology, University of Pittsburgh, Pittsburgh, PA; Pittsburgh Heart, Lung and Blood Vascular Medicine Institute, Pittsburgh, PA; Division of Cardiology, University of Pittsburgh School of Medicine, Pittsburgh, PA; College of Medicine, Alfaisal University, Riyadh, Saudi Arabia; Université Côte d’Azur, CNRS, IPMC, Valbonne, France; Division of Pulmonary, Allergy and Critical Care Medicine, University of Pittsburgh School of Medicine, Pittsburgh, PA; Department of Pathology, Boston Children’s Hospital, Boston, MA; Division of Pulmonary and Critical Care Medicine, Brigham and Women’s Hospital, Boston, MA; Division of Pulmonary and Critical Care Medicine, Department of Medicine, David Geffen School of Medicine, UCLA, Los Angeles, CA; Division of Pulmonary and Critical Care Medicine, Zuckerberg San Francisco General Hospital and Trauma Center, University of California San Francisco, San Francisco, CA

**Keywords:** pulmonary hypertension, endothelial dysfunction, pulmonary vascular changes

## Abstract

**Objective:** Pulmonary hypertension (PH) is a cardiopulmonary disease manifesting in increased pulmonary arterial pressure and right ventricular dysfunction. PH pathogenesis involves extensive pulmonary vascular remodeling precipitated, at least in part, by endothelial reprogramming. Mounting evidence points to endothelial-to-mesenchymal transition (EndMT) as an important potentiator of endothelial reprogramming in PH, yet progress in dissecting these processes remains limited.

**Approach and Results:** Lung samples from pulmonary arterial hypertension (PAH) patients and two rodent models of PH were used. Expression of the scaffolding protein ezrin-radixin-moesin-binding phosphoprotein 50 (EBP50, or NHERF1) was downregulated in PAH patient pulmonary arteries and isolated pulmonary arterial endothelial cells (PAECs), and in PH animal lung tissue and mouse isolated PAECs. In human PAECs in vitro, EBP50 was downregulated by PH-relevant stimuli, hypoxia and proinflammatory cytokine interleukin-1 beta (IL-1β). Phenocopy of EBP50 reduction in PAECs time-dependently increased expression and nuclear abundance of EndMT transcription factors Snail and Zeb1, and potentiated hypoxia-driven upregulation of Slug. Loss of EBP50 also drove expression of mesenchymal markers S100A4, fibronectin, N-cadherin, and transgelin (SM22), and inhibited cell proliferation and barrier function. In vivo studies on female EBP50^+/-^ mice demonstrated that downregulation of EBP50 exacerbated the chronic hypoxia-induced rise in RV maximum pressure.

**Conclusions:** These data identify EBP50 as a key regulator of EndMT in PH whose expression is downregulated in PH patient pulmonary endothelium and whose partial deletion exacerbates PH disease manifestations in rodents, opening doors for future therapeutic strategies to target EBP50 restoration to reverse PH.

## Introduction

Pulmonary hypertension (PH) is a multi-etiological progressive disease of heart and lung circulation defined at the 6th World Symposium on Pulmonary Hypertension (WSPH) as a rise in mean pulmonary arterial pressure (mPAP) ≥ 20mmHg ^1^. Based on common pathophysiological and hemodynamic characteristics, the WSPH guidelines distinguish five disease subgroups of PH: pulmonary arterial hypertension (PAH, = Group I PH), PH due to left heart disease (Group II), PH due to lung diseases and / or hypoxia (Group III), PH due to pulmonary artery obstructions (Group IV), and PH with unclear or multifactorial mechanisms (Group V) ^1^. In patients diagnosed with PAH, a chronic increase in pulmonary arterial pressure is associated with numerous vascular and cardiac pathologies, including right ventricular dysfunction and hypertrophy, which can ultimately be lethal ^2^. At the tissue level, PH patients present with signs of extensive pulmonary arterial remodeling and dysregulated cross-talk between pulmonary vascular cells which affects the entire arterial wall ^3^. The resulting phenotype, featuring perivascular inflammation, vascular wall thickening, rarefaction and stiffening, and luminal narrowing and occlusion ^4,5^, is thought to be initiated, at least in part, by extensive endothelial activation ^6,7^. Increased endothelial proliferation and apoptosis resistance, reduced anticoagulant properties, imbalanced production of vasoactive compounds favoring sustained vasoconstriction, metabolic dysfunction, proinflammatory shift and endothelial-to-mesenchymal transition (EndMT) are thought to disrupt the normal intracellular cross-talk within the vascular wall and promote smooth muscle hyperplasia, immune cell infiltration and sustained inflammation, ultimately resulting in a disease phenotype ^6,7^. The origins of pathological vascular changes in PH, however, remain incompletely understood.

Of many proteins with implicated functions in vascular biology, ezrin-radixin-moesin- (ERM)-binding phosphoprotein 50 (EBP50), a scaffolding PDZ protein directly interacting with adaptor proteins of the ERM family and with cytoskeletal F-actin ^8^, presents a special interest because of its role in regulating systemic smooth muscle cell (SMC) migration and proliferation ^9–11^, reactive oxygen species (ROS) production ^12^ and inflammation ^13^, all processes implicated in PH ^14,15^. Interestingly, in addition to SMCs, EBP50 was shown to control endosomal protein trafficking in systemic endothelial cells ^16^, play an important role in angiogenesis and neovascularization ^17,18^, and regulate cell division in pulmonary artery endothelial cells ^19^. However, its role in PH has never been investigated.

In recent years, as more attention has been brought to the role of pulmonary endothelium in PH pathogenesis, a number of prominent researchers in the field drew focus to significant similarities between endothelial and SMC pathophenotypes in PH and cancer ^5,7,20,27^. Interestingly, aberrant expression and subcellular localization of EBP50 has been increasingly studied in the context of various cancers ^22–24^, and several studies highlighted the role of EBP50 in regulating the activity of β-catenin ^25–27^, effector of the Wnt signaling pathway with known roles in carcinogenesis, tumor progression, and PH ^28,29^. Mechanistically, the roles of Wnt-β-catenin pathway in cancer have been linked to EndMT, a process that has recently also been implicated in PH pathogenesis by multiple research groups ^30–33^. Cells co-expressing endothelial and smooth muscle markers were found in the pulmonary vasculature of PH patients and animal models, and studies have shown that endothelial cells isolated from small pulmonary arteries explanted from PH patients and animal models exhibit signs of EndMT, including loss of endothelial markers and gain of smooth muscle markers ^32–35^. Ultimately, genetic cell reprogramming and phenotypic switching seen in EndMT result in a change in cell shape from compact cobblestone to spindle-like due to cytoskeletal remodeling and alterations in extracellular matrix composition ^31^, and potentiate an increase in endothelial monolayer permeability ^32,36^, impairing endothelial barrier function essential for the maintenance of vascular wall integrity ^7^. However, the underlying mechanisms triggering the process of EndMT in PH vessels remain unknown.

The scarcity of therapeutic interventions targeting cellular origins of PH-associated vascular remodeling and endothelial activation calls for the identification of novel molecular regulators of endothelial phenotypes in health and disease. Considering connections between EBP50 and pathways implicated in cancer and EndMT, and commonalities between cancer signaling and PH, in the current study, we sought to identify a potential role of EBP50 in PH and hypothesized that it functions as a regulator of EndMT.

## Materials and Methods

Human tissue samples were obtained through the University of Pittsburgh Tissue Processing Core, Boston Children’s Hospital, Brigham and Women’s Hospital, and University of California, Los Angeles, and approved by the University of Pittsburgh, Boston Children’s Hospital, Brigham and Women’s Hospital, and University of California, Los Angeles Institutional Review Boards. Human pulmonary artery endothelial cells from de-identified PAH patients and non-diseased subjects were obtained from the University of Pittsburgh VMI Cell Processing Core and Pulmonary Hypertension Breakthrough Initiative (PHBI) in agreement with the University of Pittsburgh and PHBI protocols. All animal experiments were approved by and conducted in accordance with the University of Pittsburgh Institutional Animal Care and Use Committee. Expanded Methods section can be found in the Data Supplement.

### Animals

For monocrotaline (MCT)-induced severe PH, 10-14 weeks old adult male Sprague-Dawley rats (Charles River Laboratories, Hudson, NY) were injected once with 60 mg/kg of monocrotaline and euthanized after 3 weeks ^37^. For chronic hypoxia-induced PH, 10-12 weeks old adult male C57BL/6 wild type mice (Taconic Biosciences, Germantown, NY and Jackson Laboratory, Bar Harbor, ME) were subjected to continuous normobaric hypoxia (10% O_2_, BioSpherix, Parish, NY) for three to four weeks (21 – 28 days). At the termination of all animal studies, lung samples were snap frozen in liquid nitrogen and stored for later analysis.

### EBP50 Heterozygous mice

7-16 weeks old adult male and female EBP50 Heterozygous mice (EBP50^+/-^, EBP50 Het), EBP50 Knockout (EBP50 KO), or wildtype (WT) controls bred on C57BL/6 background were subjected to continuous normobaric hypoxia (10% O_2_, BioSpherix, Parish, NY) for four weeks.

### Cell culture

Human pulmonary artery endothelial cells (HPAECs) were obtained from PromoCell GmbH, Germany and grown in the Endothelial Cell Growth Medium 2 (PromoCell, Germany) in the absence of heparin according to manufacturer’s protocol.

### Cell treatments

Recombinant human IL-1β was obtained from Peprotech, Inc. For hypoxia exposure, cells were cultured in temperature-humidity controlled chamber under pO_2_ = 1%, pCO_2_ = 5%, with N_2_ balance at 37°C for 48 – 72 hrs.

### EBP50 knockdown

To induce EBP50 gene knockdown, HPAECs were transiently transfected with Silencer Select EBP50 siRNA using the Lipofectamine 3000 transfection reagent according to the manufacturer’s protocol for 24 – 72 hrs.

### Quantitative real-time PCR

Levels of mRNA transcripts were quantified as previously described ^38^. TaqMan probe and Universal PCR Master Mix and SLC9A3R1, SNAI1, SNAI2, Zeb1, Zeb2, CTNNB, PECAM1, CDH5, CDH2, TAGLN, S100A4, FN1, ACTB or 18S primers (Life Technologies, Carlsbad, CA) were used for qPCR in a QuantStudio 6 Flex Real-Time PCR System (Applied Biosystems, Foster City, CA) according to the manufacturer’s protocol for 40 cycles. Relative quantification to corresponding control was obtained using the threshold cycle (Ct) method with 18S as the housekeeping gene and relative expression calculated as 2^-ΔΔCt^.

### Western blot

Western blot experiments were performed as described previously ^38^. Membranes were blocked with the Odyssey Blocking Buffer (LI-COR Biosciences, Lincoln, NE) and incubated with EBP50 (Invitrogen, Carlsbad, CA), Zeb1 (Danvers, MA), PECAM (Cell Signaling Technology, Danvers, MA), VE-cadherin (Cell Signaling Technology, Danvers, MA), N-cadherin (Cell Signaling Technology, Danvers, MA), SM22 (Abcam, United Kingdom), vinculin (Santa Cruz Biotechnology, Dallas, TX), or β-actin (Santa Cruz Biotechnology, Dallas, TX) antibodies. Membranes were then probed with fluorescence-tagged (680 or 800 nm) secondary antibodies (LI-COR Biosciences, Lincoln, NE). Digital membrane scans were obtained using the Odyssey Infrared Imaging system (LI-COR Biosciences, Lincoln, NE). Optical density of protein-of-interest bands was quantified using ImageJ software (NIH, USA) and normalized to vinculin or β-actin.

### Tissue immunofluorescence

Immunofluorescence staining was performed as previously described ^37^. Samples were blocked in 10% donkey serum and incubated with anti-EBP50 (Invitrogen, Carlsbad, CA), anti-PECAM (Abcam, United Kingdom), anti-α-SMA (Sigma Aldrich, St. Louis, MO) over night at 4°C, followed by Alexa 488-, 594- and 633-fluorophore-conjugated secondary antibodies for 1hr at room temperature, and DAPI. Images were captured using ZeissLSM780 confocal microscope. Small pulmonary vessels (<100 μm diameter) present in a given tissue section (>10 vessels / section) not associated with bronchial airways were selected for analysis. The vessel wall was defined based on α-SMA staining, and intensity of EBP50 in the vessel wall was quantified using ImageJ software (NIH, USA).

### Cells immunofluorescence

HPAECs grown on glass coverslips coated with rat tail collagen I (Gibco, Gaithersburg, MD) were fixed in 2% parafolmadehyde, permeabilized with 0.1% Triton X-100, and washed with PBS. Samples were blocked in 2% goat serum for 45 mins at room temperature and incubated with anti-Snail (R&D Systems, Minneapolis, MN), anti-Zeb1 (Invitrogen, Carlsbad, CA), anti-SM22 (Abcam, United Kingdom) and anti-PECAM (Cell Signaling Technology, Danvers, MA) antibodies, followed by 465-, Cy3- and Cy5-fluorophore-conjugated secondary antibodies. Samples were then stained for nuclei (DAPI) and affixed on slides using gelvatol mounting media. Primary delete was used as a negative control. Confocal images (15 images per treatment group) were captured using a Nikon A1 spectral confocal microscope. Intensity of staining was quantified using NIS-Elements software (Melville, NY).

### Statistics

All results are reported as mean ± SEM. For in vitro studies, where normality was not tested, normal distribution was assumed. Unpaired two-tailed Student’s t-test was used for comparison between two groups. Unpaired two-tailed Student’s t-test adjusted for the number of comparisons was used for comparison between three groups or more. To test the simultaneous effect of EBP50 and hypoxia in vitro and in vivo, we used a test of interaction from a two-way ANOVA with bootstrapping. All variables tested had passed the test for normality. For all analyses, p < 0.05 was deemed statistically significant. GraphPad Prism Software v7 and Stata 16.1 were used for data analyses.

## Results

### EBP50 expression is downregulated in the lungs of PAH patients

The handful of studies of EBP50 in vascular biology are primarily focused on systemic circulation while the role of EBP50 in pulmonary circulation remains unclear ^9,10,12,17^. Owing to the cancer-like state of endothelial cells in PH ^21^, the role of impaired redox signaling in the disease in general and ECs in particular ^39,40^, and a known association between EBP50, cancer, and PH-related processes of migration, proliferation and ROS production ^12,22–24^, we sought to determine whether EBP50 is involved in pulmonary vascular remodeling in PH. We first tested for alterations in EBP50 expression in human PAH pulmonary vasculature by comparing immunofluorescent staining of lung tissue specimens from patients with PAH to matched non-PAH controls (**Fig.1**). PAH patient demographics are detailed in **Supplement Table 1**. Using laser confocal microscopy to detect EBP50 expression, we observed a decrease in EBP50 fluorescence in and around resistance pulmonary arteries < 100 μm in diameter in PAH patients compared to controls (red pseudocolor; 0.55 ± 0.07-fold, p < 0.01, n = 4 control, 8 PAH). In addition, we predictably observed an associated increase in PAH vessel muscularization evidenced by an increased medial alpha-smooth muscle actin (α-SMA) staining (white), and a loss of continuous platelet endothelial cell adhesion molecule (PECAM) staining in the intima of PAH vessels (green), supporting a dysregulation of endothelial layer integrity and, potentially, EndMT (**Supplement Figure S1**). Importantly, colocalization of EBP50 with PECAM in control tissue and a decrease in endothelial EBP50 in PAH lungs (**Supplement Figure S2**) suggested an endothelial component of pulmonary vascular EBP50 involvement. This finding supported translational importance of EBP50 downregulation to PAH that warranted further investigation into its role in disease progression. Because of the limited availability of control lung tissues from younger patients, we were unable to fully gender- and age-match the two sample groups. Nonetheless, considering a prevalence of aging-associated degenerative processes in the lung and lung vasculature ^41,42^, our data demonstrate downregulation of vascular EBP50 expression across all ages of our PAH cohort even when compared to an older control group.

**Figure 1.**
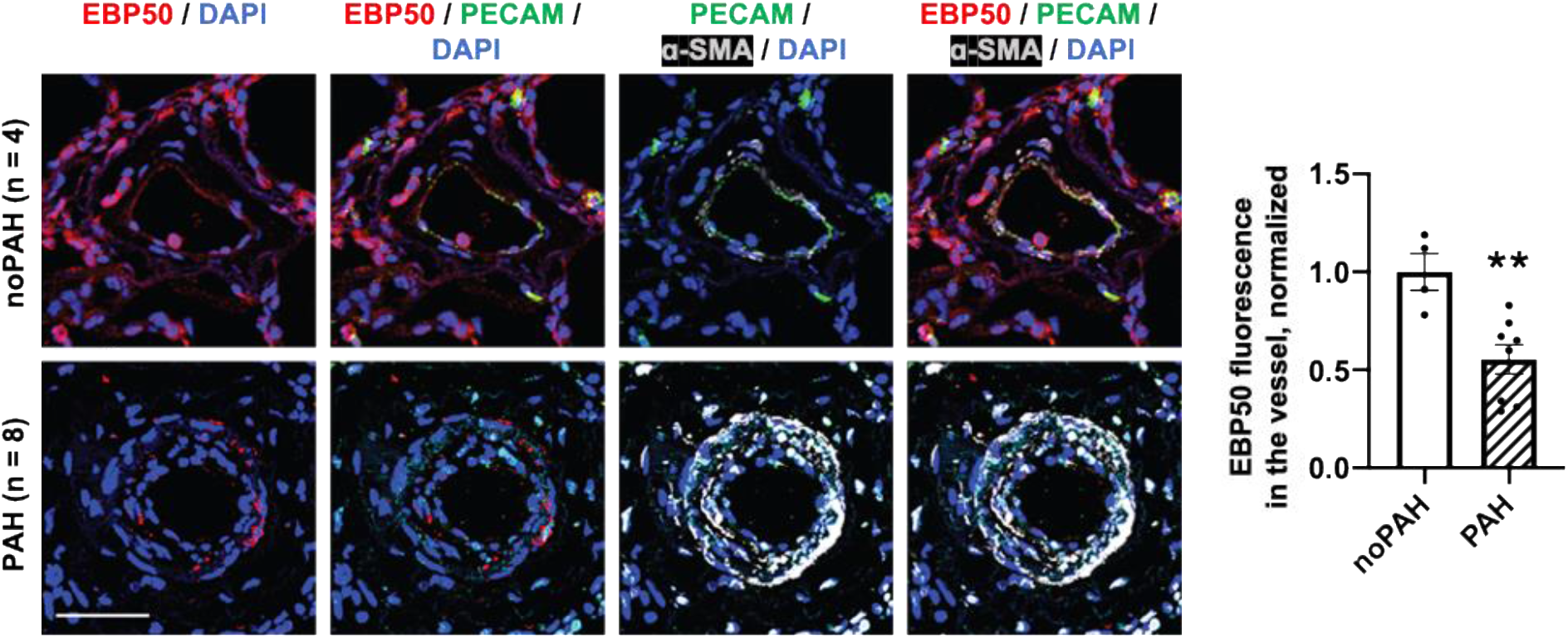
EBP50 expression is decreased in the lungs of PAH patients. Immunofluorescent staining against EBP50 (red) is decreased in PAH lung tissue vessels compared to non-PAH controls (noPAH). Staining for endothelial cell marker PECAM (green) and smooth muscle marker alpha-smooth muscle actin (α-SMA, white) reveals a loss of endothelial cell layer integrity and an increase in muscularization in PAH lung vessels (Scale bar = 50 μm). Bar graph shows quantification of the immunofluorescent signal of EBP50 per tissue per vessel and demonstrates a decrease in EBP50 signal in PAH patients relative to non-PAH controls (n = 4, control; n = 8, PAH. ** p < 0.01).

### Pulmonary EBP50 expression is downregulated in mouse and rat models of PH

In order to understand whether downregulation of EBP50 is a universal feature of PH lung tissue, we tested whether our human findings are replicated in two well-established preclinical rodent models of PH (**Fig. 2**). Immunostaining against EBP50 in PH monocrotaline (MCT)-injected rats revealed co-localization with PECAM in control tissue and a decrease in endothelial EBP50 in PH animals (**Supplement Figure S2**), and showed a decrease in pulmonary perivascular EBP50 expression (**Fig.2A, B**; 0.39 ± 0.07-fold from Vehicle, p < 0.001, n = 6), accompanied by the loss of intimal PECAM staining, an increase in medial α-SMA, as well as an increased overlap between α-SMA- and PECAM+ cells in the intima similar to changes seen in human PAH patients (**Supplement Figure S1**). In addition, EBP50 mRNA (**Fig. 2C**; 0.78 ± 0.04-fold, p < 0.05, n = 4) and protein (**Fig. 2D**; protein: 0.41 ± 0.07-fold, p < 0.05, n = 5) expression in total lung homogenates from MCT animals were decreased compared to vehicle-injected controls . Similarly, total lung EBP50 mRNA level was decreased in PH mice subjected to chronic hypoxia (**Fig. 2E**; mRNA: 0.73 ± 0.07-fold from normoxia, p < 0.05; n = 5-6). Consistent downregulation of EBP50 in mild and severe disease models in two animal species corroborated our patient findings and indicated a potential universal involvement of EBP50 in development or progression of different PH WSPH groups.

**Figure 2.**
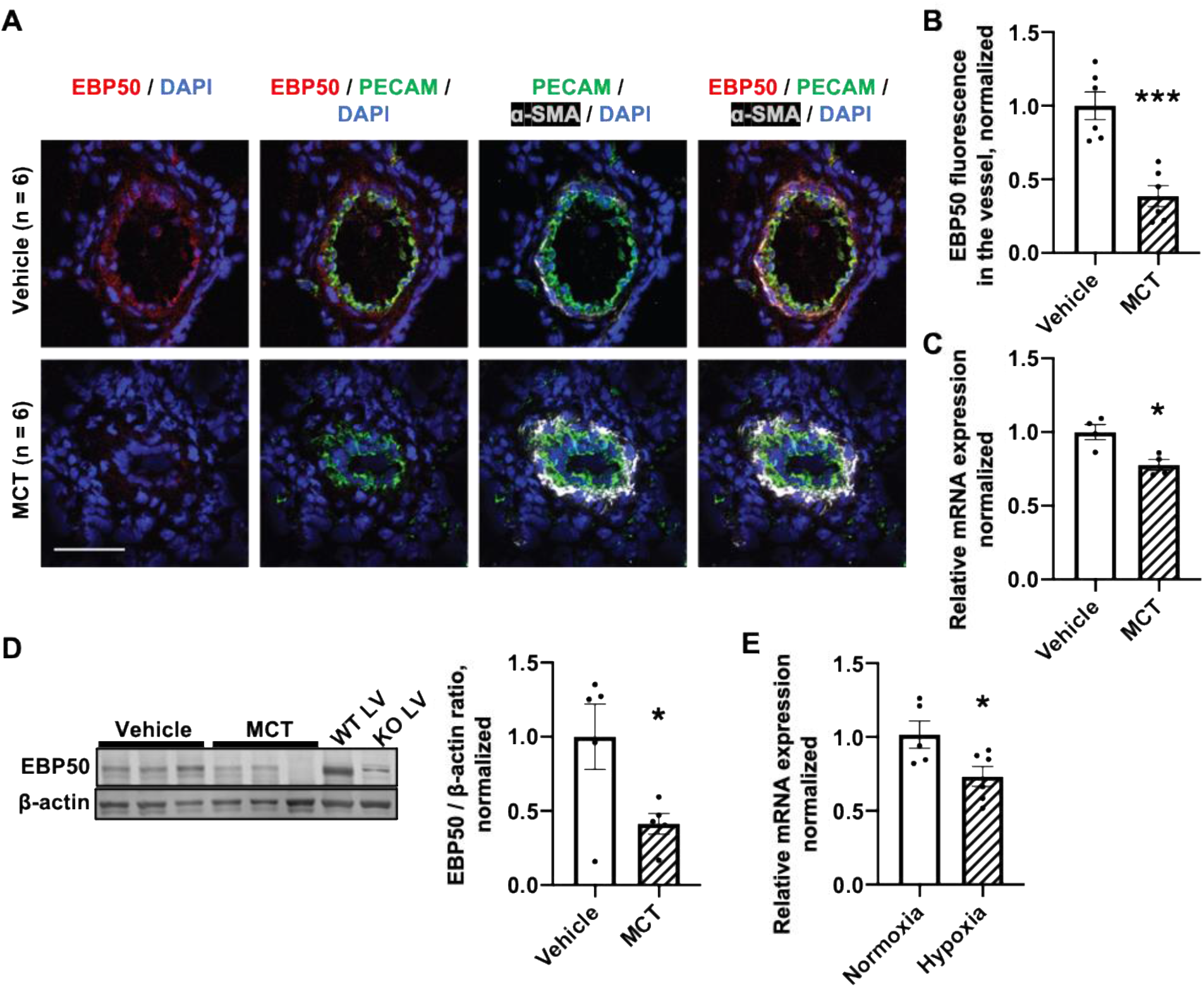
EBP50 expression is decreased in the lungs of PH rodents. A. Immunofluorescent staining against EBP50 (red) is decreased in PH lung tissue vessels of monocrotaline-treated rats (MCT) compared to Vehicle control group. Staining for PECAM (green) and α-SMA (white) reveals a decrease in endothelial marker expression in PAH vessels and an increase in vascular muscularization (Scale bar = 50 μm).
B. Quantification of the immunofluorescent signal of EBP50 per animal per vessel demonstrates a decrease in EBP50 signal in MCT rats compared to vehicle-treated controls (n = 6. *** p < 0.001).
C. EBP50 mRNA expression is decreased in the total lung homogenates from the MCT-treated PH rats compared to Vehicle-injected controls (n = 4. * p < 0.05).
D. Representative Western immunoblot (left) and densitometric quantification (right) demonstrates a decrease in EBP50 protein expression in the total lung homogenates from the MCT-treated rats compared to Vehicle controls (n = 5. * p<0.05). Left ventricle (LV) tissue from wildtype (WT) and EBP50 knockout (KO) were used as controls.
E. EBP50 mRNA expression is decreased in the total lung homogenates from the chronic hypoxia-exposed PH mice compared to normoxia controls (n = 11-14, * p < 0.05).

Relative EBP50 mRNA expression normalized to 18s or β-actin was quantified using ΔΔCt rt-PCR method. Relative EBP50 protein expression was quantified as a ratio to vinculin.

### EBP50 expression is downregulated in PAH patient- and PH mouse-derived pulmonary vascular endothelial cells

Considering our data supporting an endothelial signature of EBP50 expression, as well as the importance of endothelial cells for the maintenance of vascular homeostasis and a key role of pulmonary endothelial dysfunction in PH-associated vascular remodeling ^6,7^, we assessed whether expression of EBP50 was specifically affected in pulmonary ECs of PAH patients and PH mice. EBP50 protein levels were analyzed in endothelial cells harvested from first and second order pulmonary arteries (0.85 – 1.33 mm average diameter ^43^) of human PAH patients via immunoblotting. We used cells isolated from 6 PAH and 6 age- and sex-matched non-PAH control patients as detailed in **Supplement Table 2**. In each cohort there were 4 female and 2 male patients (2:1 female : male), consistent with the established observation of a higher incidence of PAH in females ^44,45^. To verify antibody specificity, commercially-obtained human pulmonary artery endothelial cells (HPAECs) were transfected with EBP50 siRNA for 72 hrs (knockdown efficiency at mRNA level ~ 80%, **Supplement Figure S4**), and used as a negative control. EBP50 expression was markedly downregulated in human PAH PAECs compared with non-PAH controls (**Fig. 3A**; 0.33 ± 0.05-fold from non-PAH, p < 0.0001, n = 6). These data were supported by PECAM+ pulmonary endothelial cells isolated from hypoxia-exposed mice (10% O_2_, 3 - 4 weeks) that showed reduction in EBP50 mRNA expression compared to normoxic controls (**Fig. 3B**; 0.75 ± 0.04-fold, p < 0.01; n = 11-14). Owing to the low yield of ECs from mouse lungs, protein expression analysis from these samples was not possible. Together these findings indicate that EBP50 downregulation observed in PH is, at least in part, localized to the endothelial layer of pulmonary vessels, and suggest an endothelium-specific role for EBP50 dysregulation in disease initiation or propagation.

**Figure 3.**
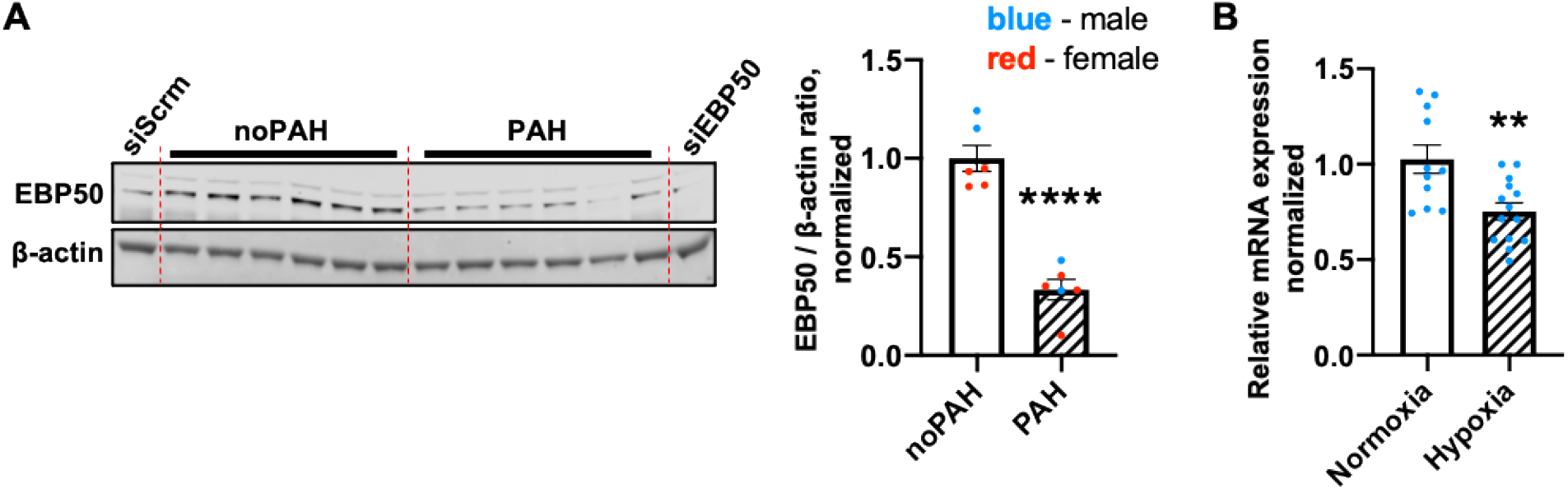
EBP50 expression is decreased in pulmonary arterial endothelial cells of PAH patients and pulmonary endothelial cells of PH chronic-hypoxia mice. A. Endothelial cells isolated from pulmonary arteries of PAH patients (n = 6) display decreased relative EBP50 protein expression compared to non-PAH donor controls (n = 6) in a protein expression analysis using Western blot (** p < 0.01). Relative EBP50 expression was quantified as a ratio to ß-actin level. Commercially obtained HPAECs transfected with EBP50 siRNA siEBP50) vs. scrambled siRNA (siScrm) for 72hrs were used as a negative control for antibody specificity (Supplement, Fig. S1).
B. Pulmonary endothelial PECAM positive cells isolated from chronic hypoxia-exposed mice (n =14) display decreased EBP50 mRNA expression compared to normoxia controls (n =11; ** p < 0.01). Relative EBP50 mRNA expression normalized to 18S was quantified using ΛΔCt rt-PCR method.

### Expression of EBP50 is downregulated in HPAECs challenged with hypoxia or PH-relevant inflammatory cytokine IL-1β

Chronic hypoxia exposure ^46^ and sustained hypoxic ^1^ and proinflammatory ^47^ environment all exacerbate and accompany PH-associated pulmonary arterial vasoocclusion and vascular remodeling. To investigate the functional relevance of EBP50 downregulation observed in pulmonary endothelial cells of PAH patients and animal PH models in vivo, we first set to determine whether PH-related stimuli induce downregulation of EBP50 in HPAECs in vitro. We used hypoxic challenge and interleukin-1 beta (IL-1β) treatment. IL-1β is an inflammatory cytokine highly upregulated in pulmonary circulation in PH patients and animals ^47,48^ and a potent inducer of transcriptional mediators of hypoxia cellular responses, hypoxia-inducible factors-1 and 2 alpha (HIF-1α and HIF-2α) ^49,50^. Both stimuli time-dependently downregulated EBP50 expression in HPAECs, albeit to a different extent. Hypoxia (1% O2) induced a ~ 40% decrease in EBP50 mRNA expression after 72 hrs, but not 48 hrs, of exposure (**Fig. 4A** & **4B**; 0.58 ± 0.07-fold at 72 hrs vs. normoxia, p < 0.01, n = 3). On the other hand, treatment with 10 ng/ml of IL-1β resulted in a marked decrease in EBP50 gene (**Fig. 4C**; mRNA: 0.51 ± 0.02-fold from untreated, p < 0.0001, n = 3) and protein (**Fig. 4D**; 0.54 ± 0.04-fold from untreated, p < 0.001, n = 3) expression as early as 48 hrs, that was sustained 72 hrs post treatment (**Fig. 4E-F**; mRNA: 0.51 ± 0.07 from untreated, p < 0.01, n = 3; protein: 0.62 ± 0.02 from untreated, p < 0.0001, n = 3). Changes in EBP50 expression after the short-term exposure to hypoxia or IL-1β are detailed in the **Supplement Figure S3**. While hypoxia exposure had no effect on EBP50 expression, IL-1β treatment induced a decrease in EBP50 mRNA expression after 3, 6, and 12 hrs (0.62 ± 0.04-fold, p < 0.05; 0.67 ± 0.05-fold, p < 0.01; 0.77 ± 0.08-fold, p < 0.05; n = 3), but not 9 hrs of exposure, suggesting a bi-phasic response.

**Figure 4.**
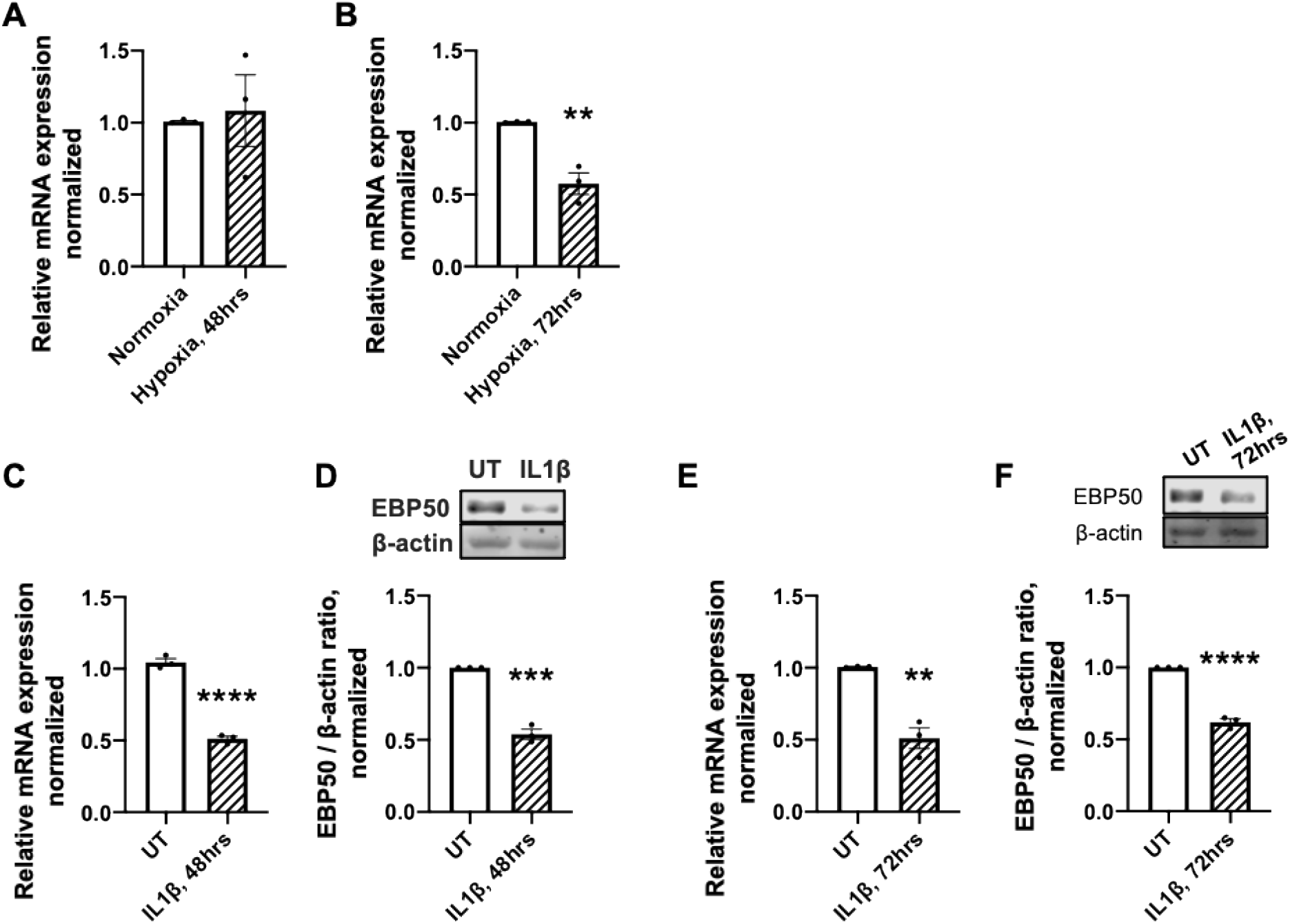
EBP50 is time-dependently downregulated by hypoxia and IL-1β treatment in HPAECs. A. 48 hrs of hypoxia exposure (pO_2_ = 1%) is not sufficient to induce stable changes in EBP50 mRNA expression (n=3).
B. EBP50 mRNA is downregulated following 72 hrs of hypoxia exposure (** - p < 0.01, n =3).
C. mRNA expression of EBP50 is downregulated following 48 hrs of 10 ng/ml IL-1 β treatment (**** - p < 0.0001, n =3).
D. Protein expression of EBP50 is downregulated following 48 hrs of 10 ng/ml IL-1β treatment (*** - p < 0.001, n =3).
E. mRNA expression of EBP50 is downregulated following 72hrs of 10 ng/ml IL-1β treatment (** - p < 0.01, n =3).
F. Protein expression of EBP50 is downregulated following 72hrs of 10 ng/ml IL-1β treatment (**** - p < 0.0001, n =3).

Relative mRNA expression was normalized to 18s and quantified using ΔΔCt rt-PCR method. Relative protein expression was calculated as a ratio to β-actin level in a respective sample.

### EBP50 knockdown upregulates EndMT transcription factors in HPAECs

While EBP50 has been functionally linked to systemic vascular SMC proliferation, migration, hypertrophy, inflammatory response, and ROS production ^9,10,12,13^, its role in the pulmonary circulation in general and in pulmonary arterial endothelial cells in particular remains elusive. Considering the known role of EBP50 as a regulator of epithelial-to-mesenchymal transition (EMT) in some cancers ^24^ and its association with β-catenin ^25–27^, one of the key drivers of EndMT, we sought to determine whether downregulation of EBP50 observed in PH models in vivo and in response to PH stimuli in vitro has a potentiating effect on EndMT in HPAECs (**Fig. 5, 6**). In phenocopy experiments, as early as 24 hrs post-transfection, EBP50 knockdown (~ 60% efficiency, **Supplement Figure S4**) significantly upregulated mRNA expression of β-catenin (mRNA, 1.43 ± 0.12-fold, p < 0.05), and EndMT transcription factors Snail, Zeb1 and Zeb2, but not Snail homolog Slug (**Fig. 5A**; 1.53 ± 0.16-fold, p < 0.05; 1.58 ± 0.05-fold, p < 0.001; 1.71 ± 0.18-fold, p < 0.05, and 1.36 ± 0.20-fold, p > 0.05, respectively, from Scramble siRNA, n = 3). Following 48 hrs of EBP50 knockdown, expression of β-catenin, Snail and Zeb1 remained upregulated (**Fig. 5B**; β-catenin: 1.67 ± 0.07-fold, p < 0.001; Snail: 1.74 ± 0.19-fold, p < 0.05; Zeb1: 1.55 ± 0.15-fold, p < 0.05 from Scramble siRNA; n = 3), and expression of Slug was increased (**Fig. 5B**; 2.05 ± 0.14-fold from Scramble siRNA, p < 0.01; n = 3). Changes in mRNA expression of Snail and Zeb1 were mirrored by increases in their nuclear abundance (**Fig. 6A,B**; Snail: 1.44 ± 0.16-fold from Scramble siRNA, p < 0.05, n = 5 after 24hrs of EBP50 knockdown and **Fig. 6C,D**; Zeb1: 1.37 ± 0.07-fold from Scramble siRNA, p < 0.01, n = 3 after 48 hrs of EBP50 knockdown) and protein expression (**Fig. 6E**; Zeb1: 1.19 ± 0.02-fold from Scramble siRNA, p < 0.01, n = 3 after 48 hrs of EBP50 knockdown), suggesting that EBP50 downregulation can drive upregulation of EndMT transcription factors and regulate Snail and Zeb1-driven transcriptional activity, as indicated by their increased nuclear abundance. Importantly, changes in expression profiles of EndMT transcription factors driven by EBP50 knockdown were similar to changes induced by IL-1β, which, in addition to its pro-PH effects, is also a potent EndMT stimulus ^51,52^ (**Supplement Figure S5**).

**Figure 5.**
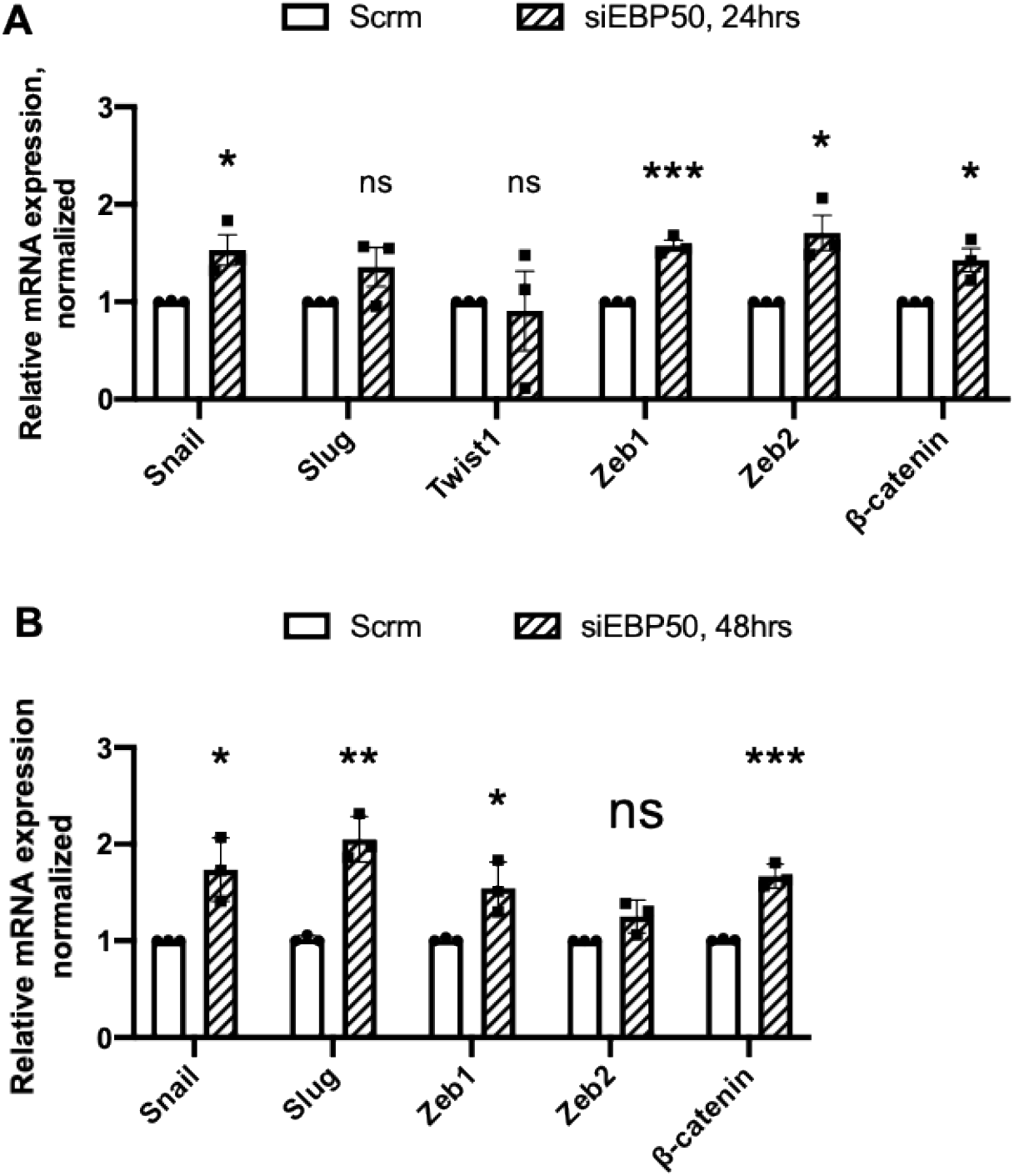
EBP50 knockdown time-dependently upregulates expression of EndMT transcription factors. A. mRNA levels of EndMT transcription factors Snail, Zeb1, Zeb2, and β-catenin are upregulated following 24 hrs of EBP50 knockdown (* - p < 0.05, ** - p < 0.01 from Scrambled siRNA, n =3).
B. mRNA levels of EndMT transcription factors Snail, Slug, Zeb1, and β-catenin are upregulated following 48 hrs of EBP50 knockdown (* - p < 0.05, ** - p < 0.01, *** - p < 0.001 vs Scrambled siRNA, n = 3).

**Figure 6.**
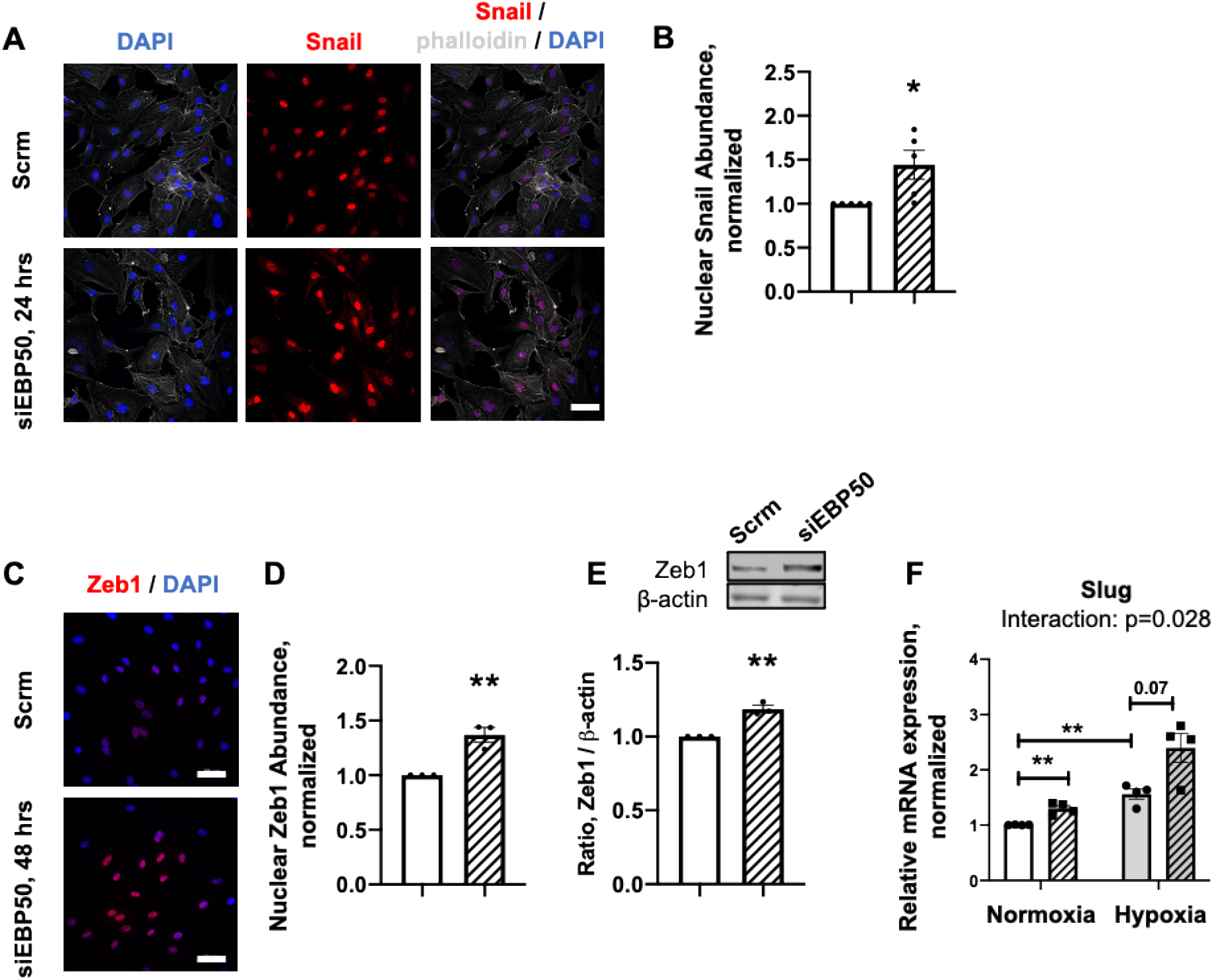
EBP50 knockdown increases nuclear abundance of EndMT transcription factors Snail and Zeb1, and increases Zeb1 protein expression. A. Nuclear immunofluorescent staining against Snail (red) is increased following 24 hrs of EBP50 knockdown (siEBP5O, 24 hrs) compared to Scrambled (Scrm) control group (Scale bar = 50 μm).
B. Nuclear abundance of Snail is increased following 24 hrs of EBP50 knockdown (* - p < 0.05 from Scrambled siRNA, n = 5).
C. Nuclear immunofluorescent staining against Zeb1 (red) is increased following 48 hrs of EBP50 knockdown (siEBP5O, 48 hrs) compared to Scrambled (Scrm) control group (Scale bar = 50 μm).
D. Nuclear abundance of Zeb1 is increased following 48 hrs of EBP50 knockdown (** - p<0.01 from Scrambled siRNA, n = 3).
E. Representative Western immunoblot (top) and densitometric quantification (bottom) of EndoMT transcription factor Zeb1 demonstrate an increase in protein expression following 48 hrs of EBP50 knockdown (** - p<0.01 from Scrambled siRNA, n = 3). Relative protein expression was calculated as a ratio to ß-actin level in a respective sample.
F. EBP50 knockdown exacerbated hypoxia-induced upregulation of Slug expression 48 hrs post-transfection. Scrm Normoxia vs. EBP50 siRNA Normoxia, ** - p<0.01; Scrm Normoxia vs. Scrm Hypoxia, ** - p<0.001; Scrm Hypoxia vs. EBP50 siRNA Hypoxia, p = 0.07; n=4. Exacerbation of the effect of hypoxia on Slug expression by EBP50 knockdown was analyzed using an interaction test from a two-way ANOVA interaction analysis with bootstrapping (p = 0.028).

mRNA normalized to 18s was quantified using ΔΔCt rt-PCR method.

We next sought to determine whether changes in expression of EndMT transcription factors driven by PH-relevant stimuli hypoxia and IL-1β are exacerbated by EBP50 dysregulation. Interestingly, while EBP50 knockdown did not have an additive effect on the expression of EndMT transcription factors stimulated by IL-1β (**Supplement Figure S6**), a combination of EBP50 knockdown and hypoxia resulted in a greater increase of Slug mRNA level following 48 hrs compared EBP50 silencing in normoxia (**Fig. 6F**). Changes in Zeb1 and Zeb2 expression displayed trends towards a differential combined effect of EBP50 silencing in hypoxia, but these were not statistically significant (p = 0.056 and p = 0.051 for Zeb1 and Zeb2, respectively). Effect of hypoxia on Snail expression was not significantly further increased by hypoxia (p = 0.91, **Supplement Figure S7**).

Together, these data indicate a role for EBP50 as a regulator of EndMT and support greater sensitivity to endothelial phenotypic changes in PH conditions where EBP50 expression is reduced.

### Knockdown of EBP50 induces gain of mesenchymal markers in HPAECs

Considering the effects of EBP50 silencing on expression and cellular localization of EndMT transcription factors, we sought to determine whether EBP50 knockdown is sufficient to induce downstream changes in endothelial and mesenchymal markers in HPAECs (**Fig. 7**). Sustained EBP50 silencing (72 hrs of knockdown, **Supplement Figure S4**) upregulated mRNA and protein for mesenchymal markers N-cadherin and SM22 (**Fig. 7B,C**; N-cadherin: mRNA, 1.77 ± 0.20-fold, p < 0.05; protein, 1.37 ± 0.15-fold, p < 0.05; SM22: mRNA, 1.89 ± 0.10-fold, p < 0.01; protein, 1.72 ± 0.08-fold, p < 0.001; compared to Scramble-transfected controls, n = 3-4), and induced expression of fibrotic mesenchymal markers S100 Calcium Binding Protein A4 (**Fig. 7B**; S100A4; mRNA, 1.66 ± 0.10-fold, p < 0.01; n = 3) and fibronectin (**Fig. 7B**; FN1; mRNA, 1.46 ± 0.18-fold, p < 0.05; n = 4). However, EBP50 knockdown did not significantly affect expression levels of endothelial markers PECAM (**Fig. 7A,B,D**; mRNA, 1.53 ± 0.41-fold; protein, 1.06 ± 0.12-fold; n = 3) and vascular endothelial cadherin (**Fig. 7A,B,D**; VE-cadherin; mRNA, 1.35 ± 0.32-fold; protein, 0.86 ± 0.07-fold; n = 3). Notably, EBP50 knockdown closely recapitulated IL-1β induction of mesenchymal marker expression N-cadherin and SM22 for the same timepoint at the mRNA and protein level, but not the IL-1β reduction of endothelial markers PECAM and VE-cadherin (**Supplement Figure S8**). These data support that EBP50 functions as a regulator of adoption of mesenchymal characteristics typical of late-stage EndMT, while having minimal effect on endothelial markers.

**Figure 7.**
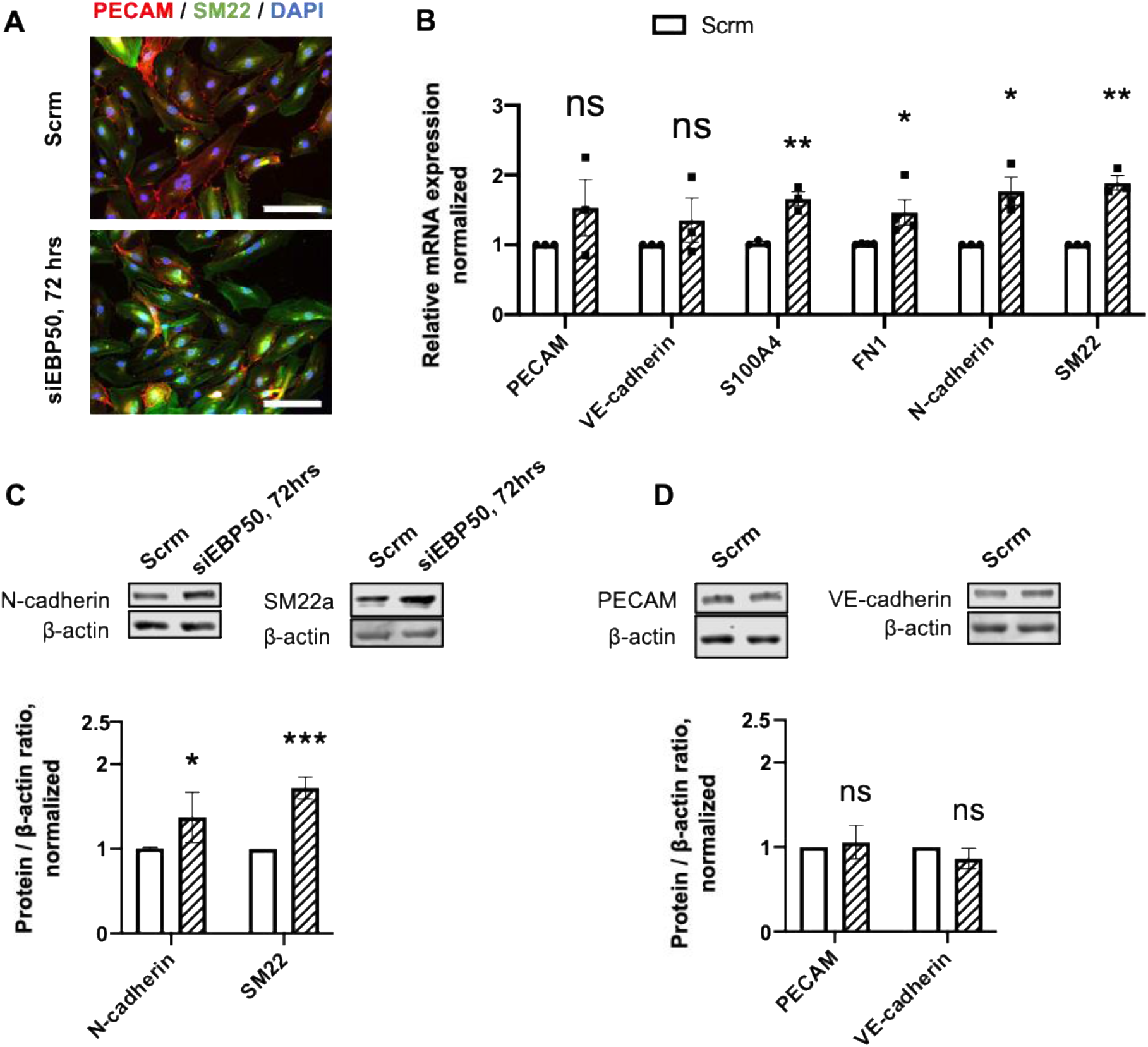
EBP50 knockdown promotes gain of mesenchymal markers by the pulmonary endothelial cells. A. Immunofluorescent staining against PECAM (red), SM22 (green) in HPAECs following 72 hrs of EBP50 knockdown (siEBP50, 72 hrs) compared to Scrambled (Scrm) control group (Scale bar = 200 μm).
B. mRNA expression of endothelial markers PECAM and VE-cadherin remains unchanged, while mesenchymal markers S100A4, fibronectin, N-cadherin and SM22 are upregulated following 72 hrs of EBP50 knockdown (* - p < 0.05, ** - p < 0.01 from Scrambled siRNA, n = 3-4). mRNA normalized to 18s was quantified using ΔΔCt rt-PCR method.
C. Representative Western immunoblots (left) and densitometric quantification (right) of N- cadherin and SM22 demonstrate an increase in protein expression following 72 hrs of EBP50 knockdown (* - p<0.05, ** - p<0.01 from Scrambled siRNA, n = 3-4). Relative protein expression was calculated as a ratio to β-actin level in a respective sample.
D. Representative Western immunoblots (left) and densitometric quantification of PECAM and VE-cadherin show no difference in protein expression following 72 hrs of EBP50 knockdown (* - p<0.05, ** - p<0.01 from Scrambled siRNA, n = 3). Relative protein expression was calculated as a ratio to β-actin level in a respective sample.

### EBP50 knockdown impairs pulmonary endothelial proliferation

While increased endothelial proliferation is recognized as a feature of PH-associated vascular remodeling ^38,53^, studies of the impact of EndMT on proliferation are not in universal agreement. While most studies link EndMT to increased cell proliferation ^34,54^, others report contrasting findings and argue that unstimulated HPAECs exhibit a greater increase in cell number compared to HPAECs treated with pro-EndMT cytokines, such as tumor necrosis factor alpha (TNFα), transforming growth factor-beta (TGF-β), and IL-1β^32^. Combined with increased permeability, disordered endothelial proliferation impairs the restoration of endothelial monolayer integrity and barrier function in response to damage, and contributes to exacerbated vascular remodeling ^32^. To test whether PH-relevant downregulation of EBP50 is associated with alterations in endothelial proliferation in HPAECs transfected with EBP50 siRNA, we quantified HPAECs proliferation using fluorescence-activated cell sorting (FACS) analysis and DNA replication and repair by monitoring protein expression of DNA polymerase processivity factor proliferating cell nuclear antigen (PCNA). Extent of *de novo* DNA biosynthesis preceding cell division was decreased in EBP50 siRNA-transfected HPAECs by ~ 30% (% proliferating cells: 19.16 ± 0.49 vs. 13.21 ± 1.13 in Scrambled siRNA-transfected vs. EBP50 siRNA-transfected HPAECs, p < 0.01, n = 3, **Fig. 8A,B**), as indicated by a lower rate of thymidine analog EdU incorporation during the S phase of the cell cycle 24 hrs post-transfection. Moreover, EBP50 knockdown produced stable downregulation of PCNA at 48 and 72 hrs of gene silencing, suggesting impairment of DNA replication or increased DNA damage (0.71 ± 0.07-fold, p < 0.05 and 0.57 ± 0.02-fold, p < 0.0001 from Scrambled siRNA, respectively; n = 3, **Fig. 8 C,D**). EBP50 knockdown did not appear to affect apoptotic cell death, as indicated by the lack of a difference in caspase 3/7 cleavage between Scrambled and EBP50 siRNA-transfected HPAECs (**Supplement Figure S9**).

**Figure 8.**
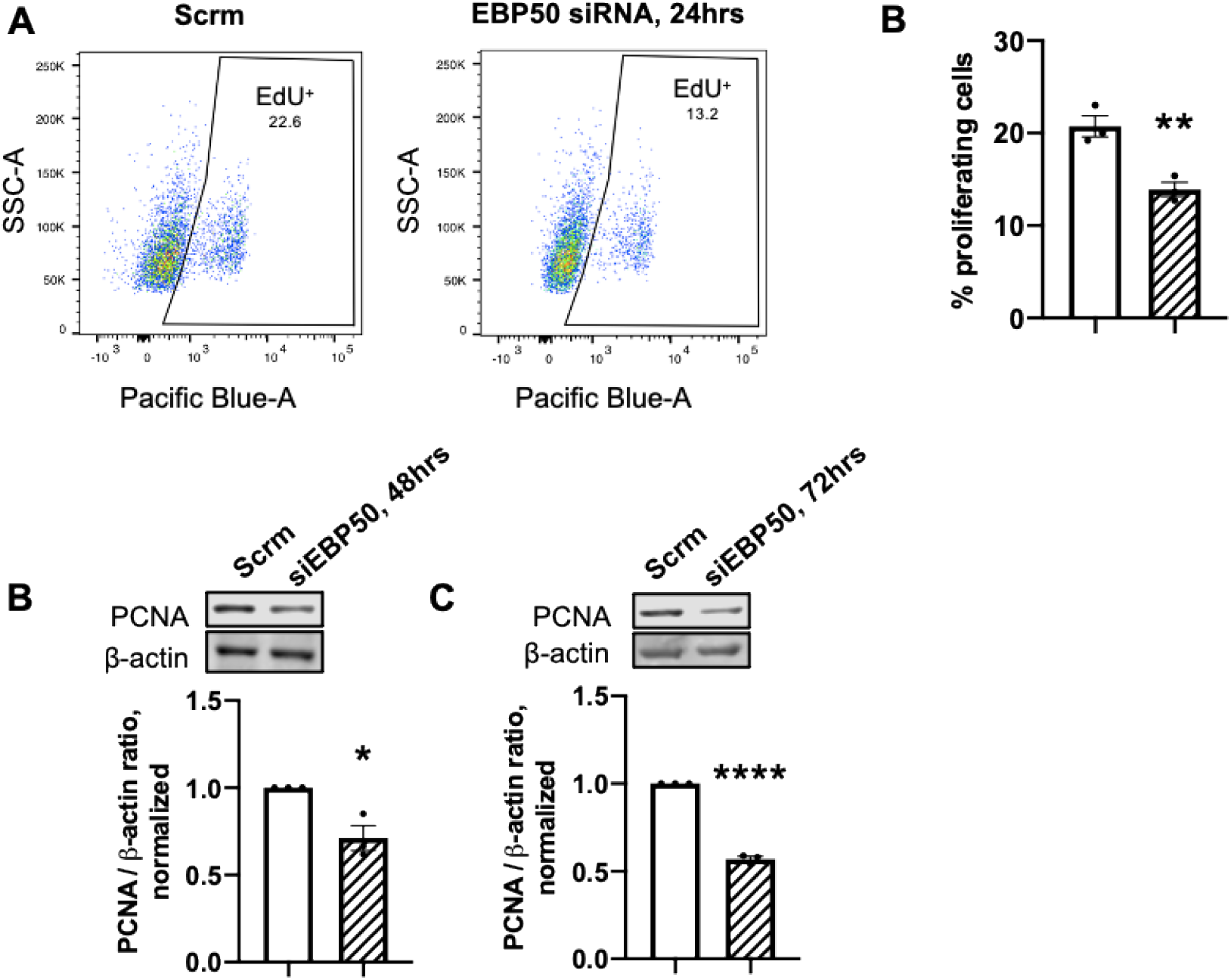
EBP50 knockdown decreases pulmonary endothelial cell proliferation and decreases baseline resistance of endothelial monolayer. A. DNA incorporation of a thymidine analog EdU measured by flow cytometry.
B. EdU incorporation is decreased in cells transfected with EBP50 siRNA for 24 hrs (** - p < 0.01 from Scrambled siRNA, n =3). B,C. Representative Western immunoblots (top) and densitometric quantifications (bottom) of proliferating cell nuclear antigen (PCNA) demonstrate a decrease in protein expression following 48 and 72 hrs of EBP50 knockdown compared to Scrambled control (* - p < 0.05, **** - p < 0.0001, n =3).

### Knockdown of EBP50 impairs endothelial barrier function and predisposes to increased endothelial monolayer permeability

Impaired endothelial barrier function and increased endothelial monolayer permeability are associated with PH ^55–57^, and EndMT ^32,36^. To determine whether EBP50 regulates endothelial permeability and drives vascular manifestations of EndMT in PH, transendothelial electrical resistance (TEER) of EBP50 siRNA-transfected HPAEC monolayer was assessed in real-time using Electric Cell-substrate Impedance Sensing (ECIS) technology ^58^ (**Fig. 9A**). In response to low frequency applied current (4000 Hz), baseline resistance of HPAECs transfected with EBP50 for 24 hrs was lower than that of Scrambled-transfected HPAECs (870.1 ± 24.1 vs. 773.2 ± 24.16 Ohm for Scrambled-vs. EBP50 siRNA-transfected cells, p < 0.05, n = 3), indicating that disruption of EBP50 has a negative effect on endothelial barrier function. However, co-stimulation of transfected cells with IL-1β did not result in a further impairment of endothelial barrier function (**Supplement Figure S10**), in line with our findings in **Supplement Figure S8** suggesting that EBP50 may not exacerbate IL-1β effects under these experimental conditions.

**Figure 9.**
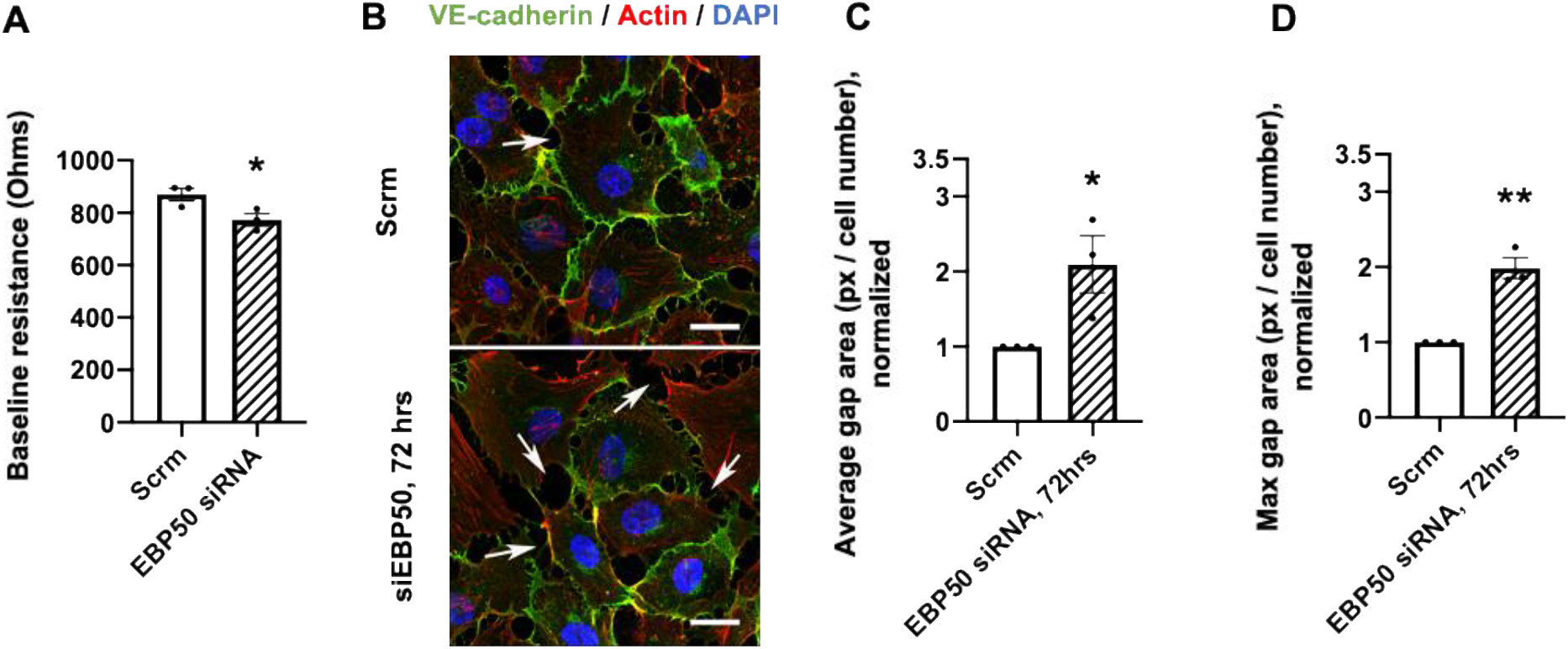
EBP50 knockdown decreases baseline resistance and increases intercellular gap size in HPAECs monolayers. A. Baseline resistance of the HPAECs monolayer transfected with EBP50 siRNA for 24 hrs is decreased compared to Scramble-transfected, indicating increased endothelial permeability (* - p < 0.05, n = 3).
B. Representative image of intercellular gap areas in Scrm- and EBP50 siRNA-transfected endothelial monolayers after 72hrs of transfection. Scale bar = 20uM.
C. Quantification of the average intercellular gap areas in Scrm- and EBP50 siRNA-transfected endothelial monolayers after 72hrs of transfection across three independent experiments (* - p < 0.05, n = 3).
D. Quantification of the maximum intercellular gap areas in Scrm- and EBP50 siRNA-transfected endothelial monolayers after 72hrs of transfection across three independent experiments (** - p < 0.01, n = 3).

Furthermore, we used immunofluorescent staining of Scrambled or EBP50 siRNA-transfected human pulmonary arterial endothelial cells for critical junction proteins VE-cadherin and ZO-1 to investigate whether EBP50 downregulation disrupts tight and adherens junctions. We then performed a quantitative image analysis calculating the ratio of abundance of ZO-1 and VE-cadherin and vice versa only in regions where both appear. Our analysis concluded that subcellular localization or relative abundance of critical cell-cell junctions proteins was not altered with EBP50 siRNA (**Supplement Figure S11**).

To provide an additional proof for the effect of EBP50 knockdown on endothelial monolayer resistance we quantified the size (in pixels) of intercellular gaps in HPAECs transfected with Scrambled and EBP50 siRNA (**Fig. 9B-D**). Our quantitative analysis indicated that cells transfected with EBP50 siRNA have a looser monolayer structure, where average and maximum cell-cell gaps areas were increased by ~50% (2.11 ± 0.38-fold, p < 0.05, and 1.99 ± 0.14-fold, p < 0.01, from Scrambled for average and maximum gap area, respectively; n = 3).

### Genetic attenuation of EBP50 in female mice exacerbates chronic hypoxia hemodynamic outcomes in vivo

Following in vitro studies indicating that EBP50 plays a role in regulating endothelial cell reprogramming and function, we sought to determine whether genetic downregulation of EBP50 exacerbates chronic hypoxia-induced PH in vivo. For these studies, since pulmonary EBP50 expression in human PAH tissue and different in vivo preclinical PH models is downregulated by approximately 50% but not entirely abolished compared to non-PH controls (**Fig. 1-2**), we chose to utilize EBP50 heterozygous (EBP50 Het, EBP50^+/-^) mice (**Fig. 10, Supplement Figure S12, 13**). However, we also looked into *in vivo* RV function parameters of EBP50 knockout (EBP50 KO) mice to test the effect of complete abolishment of EBP50 on hemodynamic manifestations of PH. Genetic interruption of EBP50 in female mice did not alter baseline RV maximum pressure (**Fig. 10A**; RVmaxP; WT Normoxia, 24.94 ± 1.34 mmHg vs. Het Normoxia, 25.59 ± 1.34 mmHg vs. KO Normoxia, 25.82 ± 1.19 mmHg, n = 6-9) or RV hypertrophy (**Fig. 10 B, C**; RV / body weight ratio: WT Normoxia, 0.88 ± 0.04 g/g vs. Het Normoxia, 0.81 ± 0.03 g/g vs. KO Normoxia, 0.94 ± 0.03 g/g; Fulton index [RV / (LV + Septum) weight]: WT Normoxia, 0.27 ± 0.01 vs. Het Normoxia, 0.25 ± 0.01 vs. KO Normoxia, 0.27 ± 0.01; n = 6-10). In contrast, there was a further rise in hypoxia-induced RVmaxP in EBP50 Het female mice subjected to hypoxia compared to hypoxia-exposed WT controls (**Fig. 10A**; WT Hypoxia: 35.56 ± 1.30 mmHg, vs. Het Hypoxia: 41.97 ± 0.77 mmHg, p < 0.01, n = 6-9). This increase was surprisingly not associated with an exacerbation of hypoxia-induced RV hypertrophy (**Fig. 10 B, C**; RV / body weight ratio: WT Hypoxia: 1.45 ± 0.05 gg, vs. Het Hypoxia: 1.31 ± 0.05 g/g; Fulton Index: WT Hypoxia: 0.44 ± 0.03, vs. Het Hypoxia: 0.39 ± 0.01 mg/mm; n = 8-11), suggesting that, in the hypoxia mouse PH model, functional effect of EBP50 loss may be initially manifested in the vasculature rather than the heart, which appeared to be adapted to the ~ 6.5 mmHg additional pressure overload. Interestingly, EBP50 knockout did not have an added effect on hypoxia-induced increase in RV maximum pressure (33.78 ± 2.43 mmHg, n = 11) or RV hypertrophy (RV / body weight ratio: 1.41 ± 0.09 g/g; Fulton index: 0.38 ± 0.02, n = 11) compared to hypoxia-exposed WT controls. Lastly, the RV contractile index (RVCI, max *dP* / d*t* normalized for RVmaxP pressure, 1 / sec) was not affected by hypoxia exposure and was stable in hypoxia-exposed Hets and KOs compared to WT controls (**Fig. 10D**). Interestingly, hypoxia-exposed male EBP50 Het mice did not demonstrate an exacerbated hypoxia-induced RVmaxP (**Supplement Figure S13**; WT Hypoxia, 35.94 ± 1.78 mmHg vs. Het Hypoxia, 35.54 ± 2.55 mmHg, n = 6), nor RV hypertrophy (RV / body weight ratio: WT Hypoxia, 1.19 ± 0.05 g/g vs. Het Hypoxia, 1.33 ± 0.07 g/g; Fulton Index: WT Hypoxia, 0.34 ± 0.02 vs Het Hypoxia, 0.36 ± 0.01, n = 6) compared to WT controls, supporting a potentially sex-specific effect of EBP50 interruption on the pulmonary vasculature. This may be related to estrogen receptor signaling and could offer insight to the prevalence of the human disease in females compared to males. Further, LV parameters (LV maximum pressure and LV hypertrophy) stayed mostly stable across experimental groups in both males and females, suggesting that the left heart was not affected (**Supplement Figure S14**). Collectively, the in vivo results closely align with our in vitro data showing that downregulation of EBP50 potentiates effects of hypoxia-driven signaling processes implicated in PH, such as EndMT.

**Figure 10.**
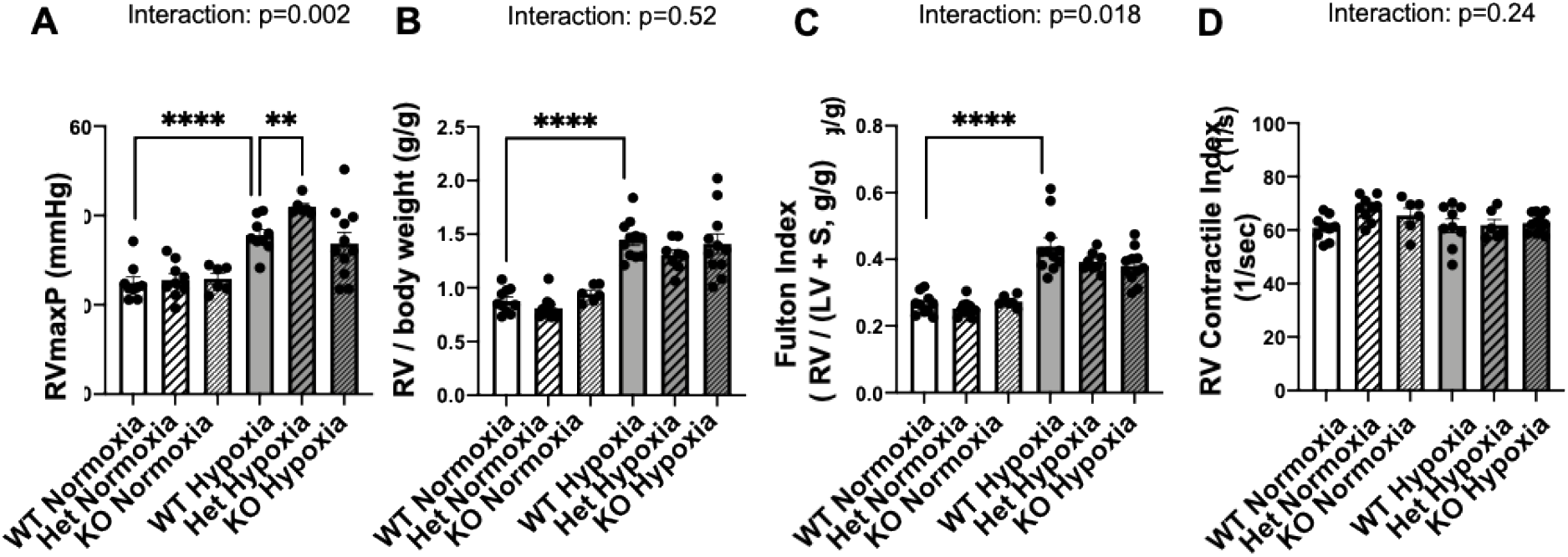
When subjected to hypoxia, EBP50 heterozygous mice exhibit an exacerbated RVmaxP compared to wild-type controls, but do not exhibit a higher degree of RV hypertrophy or changes in the RV contractile index. For a period of chronic hypoxia (4 weeks) treatment, experimental animals were placed in an environment with FiO_2_ = 10%. At the end of the treatment period, pressure-volume loop measurements were acquired through high-fidelity admittance catheter and hemodynamic parameters were calculated. Exacerbation of the effect of hypoxia on hemodynamic measures by partial EBP50 loss was analyzed using an interaction test from a two-way ANOVA interaction analysis with bootstrapping

A. Hypoxia-induced elevation of the RV maximum pressure was more pronounced in EBP50 heterozygous female mice compared to wild-type controls.
B. EBP50 Het Hypoxia mice did not exhibit an exacerbated RV hypertrophy calculated as a ratio of RV weight (g) to body weight (g) compared to WT Hypoxia group.
C. EBP50 Het Hypoxia mice did not exhibit an exacerbated RV hypertrophy (increased Fulton Index) compared to WT Hypoxia group.
D. EBP50 Het Hypoxia mice did not exhibit an altered RV Contractile Index compared to other experimental groups.

N = 6-11. ** - p <0.01, **** - p < 0.0001.

## Discussion

Despite significant progress in our understanding of molecular mechanisms driving vascular remodeling in PH, knowledge about molecular players that regulate key endothelial phenotypes associated with PH, such as EndMT, remains limited. In this study, we for the first time showed that downregulation of a scaffolding PDZ protein, EBP50, in the pulmonary arteries and pulmonary arterial endothelial cells is a shared feature between human PAH and animal models of PH. We identified disruption of EBP50 as a driver of EndMT in cultured HPAECs and demonstrated that EBP50 plays an important role in regulating cell proliferation and maintenance of endothelial barrier function. Functional importance of EBP50 downregulation was established in genetic female EBP50 heterozygous mice, which demonstrated an exacerbation of hemodynamic manifestations of PH in response to chronic hypoxia exposure compared to wild-type animals. These data identify a novel player in PH pathogenesis and offer mechanistic insights into the regulation of PH-associated EndMT.

Studies on scaffolding proteins in PH have been limited. Since about a decade ago, a handful of studies identified mutations of cytosolic and mitochondrial scaffolding proteins, such as caveolin-1 and NFU1 iron-sulfur cluster scaffold, in PH and connected them to PH-associated vascular remodeling and disease progression ^59–61^. While interest in other scaffolding proteins in PH research may have partially subsided, those proteins may still offer valuable knowledge of this devastating disease. Owing to their ability to have multiple binding partners, scaffolding proteins can exhibit cell type-specific functions and serve as cellular “rheostats” to precisely regulate distinct molecular cascades and offer a unique opportunity for targeted interruption of protein-protein interactions ^62^. In our work we expand on these studies to identify functional downregulation of the PDZ scaffolding protein EBP50 in the pulmonary arteries, whose connection to PH and PAH, to our knowledge, was not previously reported.

Indicative of its potential significance, EBP50 expression was downregulated in both animal models of PH used in the study, the mild chronic-hypoxia and severe MCT PH. Critically, we demonstrate that downregulation of EBP50 is also featured in the pulmonary arteries of PAH patients, which underscores translational importance of this finding and hints at the potential of EBP50 restoration for the treatment of PAH. Moreover, a loss of EBP50 appeared to be shared between PH endothelial cells of both humans and animals and was recapitulated in HPAECs challenged with PH-relevant stimuli, hypoxia and IL-1β. It is worth noting that in different tissues assessed by Western blot we observed multiple bands that appeared to correspond to EBP50 protein. This observation is consistent with previous studies ^63^, which identified proteolytic variants of EBP50 in total fractions and membranes from mouse kidney cortex and intestine. Moreover, a number of phosphorylation sites have been identified for EBP50 protein, and phosphorylation patterns of these residues appear asynchronous and under control of mechanisms that are incompletely understood. This necessitated use of multiple controls such as EBP50 siRNA silenced HPAEC lysates and EBP50 KO mouse heart tissue in our Western blot analyses.

Although our understanding of the signaling mechanisms contributing to PH-associated vascular remodeling is expanding continuously, much remains to be discovered about the role of endothelial reprograming and EndMT in disease progression. EndMT is a highly dynamic process, characterized by hard-to-define initiation, progression, and check-points, which pose significant challenges for studying it in vivo. To this date, little is known about where along the spectrum of PH initiation and progression EndMT manifests itself most prominently. However, the knowledge amassed in the last decade points at the importance of this process in the early events leading to a dysregulation of vascular homeostasis and impaired intercellular cross-talk within the vascular wall ^34,35^. In this work, we identified the functional role of PH-associated downregulation of EBP50 in driving EndMT in HPAECs, demonstrated by a time-dependent upregulation of EndMT transcription factors, Snail, Slug and Zeb1, and a gain of mesenchymal markers N-cadherin, SM22, S100A4, and fibronectin in the phenocopy experiments in human primary endothelial cells. However, unlike other known EndMT inducers, such as IL-1β, reduction in EBP50 alone was not sufficient to fully drive complete EndMT by itself, evidenced by a lack of added effect on levels of endothelial markers, but instead had a potentiating effect on hypoxia-induced activation of EndMT transcription factors. This observation deserves further investigation and can potentially explain our in vivo findings in female EBP50^+/-^ mice, which develop a more severe PH phenotype in response to chronic hypoxia compared to WT mice.

Consistent with earlier reports linking EndMT to impaired barrier function ^32,36^, we demonstrated that the knockdown of EBP50 increases HPAEC monolayer permeability. Combined with our identification of a role for EBP50 downregulation as an inhibitor of endothelial proliferation, these results further argue that a decreased level of EBP50, such as seen in PH vasculature, exacerbates vascular remodeling by impairing the restoration of endothelial monolayer integrity and barrier function in response to damage.

Hemodynamic analysis supports our findings of the importance of EBP50 in the pulmonary vasculature. Despite an exacerbated RVmaxP, female EBP50 Het mice in hypoxia did not develop an additional increase in RV hypertrophy or a change in the RV contractility index, indicating that the effects of EBP50 dysregulation may, in the early and somewhat moderate severity disease stages, primarily affect pulmonary arteries, rather than the heart, which may be well-adapted in the mouse hypoxia model. Interestingly, male EBP50 Het mice did not display either an exacerbated RVmaxP or RV hypertrophy after a chronic hypoxia exposure. This observation suggests that the effects of EBP50 may be closely tied to sex hormone signaling, which is consistent with the long-known role of estrogen receptor in regulating EBP50 in cancer cells ^64^ and linking the estrogen response in cancer to various cytoskeletal and cell signaling pathways ^65^. Further investigation is needed to tease out this relationship. In addition, to test for a specific role of EBP50 downregulation in the endothelium we are planning to utilize EC-specific EBP50 Het mice.

Despite the known associations between PH and a whole lung expression of EndMT transcription factors ^33,35,66,67^, the precise role of EndMT in PH remains inadequately understood. Indeed, complexities accompanying the detection of EndMT in vivo may explain our inability to detect consistent, statistically significant upregulation of EndMT markers in isolated PAH patient ECs or PECAM-positive cells from hypoxia-exposed mice (data not shown). To address this limitation in our future studies, we will consider utilizing a recently described method of fibrin gel implantation ^68^ as a way of detecting EndMT in vivo.” This too is an active area of focus in our laboratory.

The work herein enhances our understanding of the role of ERM-binding proteins in vascular biology in general and PH in particular, and maps out potential avenues for development of therapeutics specifically targeting endothelial cells in PH. Despite the fact that PDZ scaffolding proteins are abundantly expressed in mammals, and that 200 – 300 PDZ proteins are encoded in the human genome ^69^, the number of studies concerning PDZ proteins in PH is low. PDZ proteins regulate a wide array of cellular functions, controlling receptor trafficking in synapses ^70^ and regulating drug transporters ^71^ and ion exchangers ^72^. Yet, to our knowledge, only a handful of studies connected alterations of expression of PDZ proteins to PH, implicating only PDZ And LIM Domain Protein 5, PDLIM5 ^73,74^, and Shroom ^75^. Here we expand on this literature and present data that support the development of therapeutics aiding the restoration of normal EBP50 expression in pulmonary arterial endothelial cells and open the door to more in-depth investigation of the link between EBP50 loss, EndMT and PH progression in vivo. Our work sheds some light on these mechanisms and introduces EBP50 as a potential upstream regulator. Future work is expected to elucidate the mechanistic connection between EBP50 downregulation and vascular remodeling in PH and will pave the way for novel avenues for therapeutic strategies for this devastating disease.

## List of Abbreviations

EBP50: Ezrin-radixin-moesin-binding phosphoprotein 50
ECIS: Electric Cell-substrate Impedance Sensing
EndMT: Endothelial-to-mesenchymal transition
HPAEC: Human pulmonary artery endothelial cell
IL-1β: Interleukin-1 beta
LV: Left ventricle
PA: Pulmonary artery
PAEC: Pulmonary artery endothelial cell
PAH: Pulmonary arterial hypertension
PCNA: DNA polymerase processivity factor proliferating cell nuclear antigen
PH: Pulmonary hypertension
RV: Right ventricle
RVmaxP: Right ventricle maximum pressure
S100A4: S100 Calcium Binding Protein A4
SM22: Transgelin
SMC: Smooth muscle cell
TEER: Transendothelial electrical resistance

## Acknowledgements

We would like to thank Drs. Claudette St. Croix, Jonathan Florentin and Partha Dutta for their help with generating preliminary data for this manuscript. The graphic abstract in this manuscript was created with BioRender.com.

## Highlights

- Expression of scaffolding PDZ protein EBP50 is downregulated in in PAH patients and PH and PAH rodent ECs as well as in vitro in human primary ECs treated with PH-related stimuli (hypoxia and IL-1β).
- Downregulation of EBP50 promotes EndMT in pulmonary ECs and impairs cell proliferation and barrier function, processes reflective of PH cellular phenotypes.
- EBP50 Heterozygous mice subjected to chronic hypoxia display worsened hemodynamic outcomes than WT mice supporting the importance of EBP50 modulation as key to PH pathogenesis.

