## Supplemental Figures for "The Role of EBP50 in Regulating Endothelial-To-Mesenchymal Transition in Pulmonary Hypertension"

### The Role of Ezrin-Radixin-Moesin-Binding Phosphoprotein 50 in Regulating Endothelial-To-Mesenchymal Transition in Pulmonary Hypertension

Anastasia Gorelova et al.

#### Supplemental Materials

| Group | ID # | Sex | Age | PH group | Type PH |
|---|---|---|---|---|---|
| **control** | 1 | Male | 72 | N/A | N/A |
| | 2 | Female | 69 | N/A | N/A |
| | 3 | Male | 68 | N/A | N/A |
| | 4 | Male | 65 | N/A | N/A |
| **PAH** | 1 | Female | 34 | Group 1 PH | Idiopathic |
| | 2 | Female | 64 | Group 1 PH | Idiopathic |
| | 3 | Female | 68 | Group 1 PH | Scleroderma |
| | 4 | Male | 12 | Group 1 PH | Familial (BMPR2 mutation) |
| | 5 | Male | 16 | Group 1 PH | Idiopathic |
| | 6 | Male | 1 | Group 1 PH | Trisomy 21 |
| | 7 | Male | 19 | Group 1 PH | Idiopathic |
| | 8 | Female | 42 | Group 1 PH | Scleroderma |

**Supplement Table 1. Demographic and clinical characteristics of non-PAH control and PAH patient population used for immunofluorescent staining.**

| Group | ID # | Sex | Age | PH group | Type PH | mPAP, mmHg | PCWP, mmHg | PVR, WU |
|---|---|---|---|---|---|---|---|---|
| **control** | 1 | female | 53 | N/A | N/A | N/A | N/A | N/A |
| | 2 | male | 25 | N/A | N/A | N/A | N/A | N/A |
| | 3 | female | 34 | N/A | N/A | N/A | N/A | N/A |
| | 4 | male | 24 | N/A | N/A | N/A | N/A | N/A |
| | 5 | female | 36 | N/A | N/A | N/A | N/A | N/A |
| | 6 | female | 50 | N/A | N/A | N/A | N/A | N/A |
| **PAH** | 7 | male | 53 | group 1 PH | idiopathic | 56 | 5 | 3.86 |
| | 8 | male | 21 | group 1 PH | idiopatnic | 69 | 12 | 19.74 |
| | 9 | female | 62 | group 1 PH | familial | 48 | 8 | 8 |
| | 10 | female | 29 | group 1 PH | idiopathic | 110 | 19 | 6.29 |
| | 11 | female | 16 | group 1 PH | idiopathic | N/A | 10 | N/A |
| | 12 | female | 40 | group 1 PH | idiopathic | N/A | N/A | N/A |

**Supplement Table 2. Demographic characteristics and hemodynamic parameters of non-PAH control and PAH patient populations used for pulmonary endothelial cell isolation.**

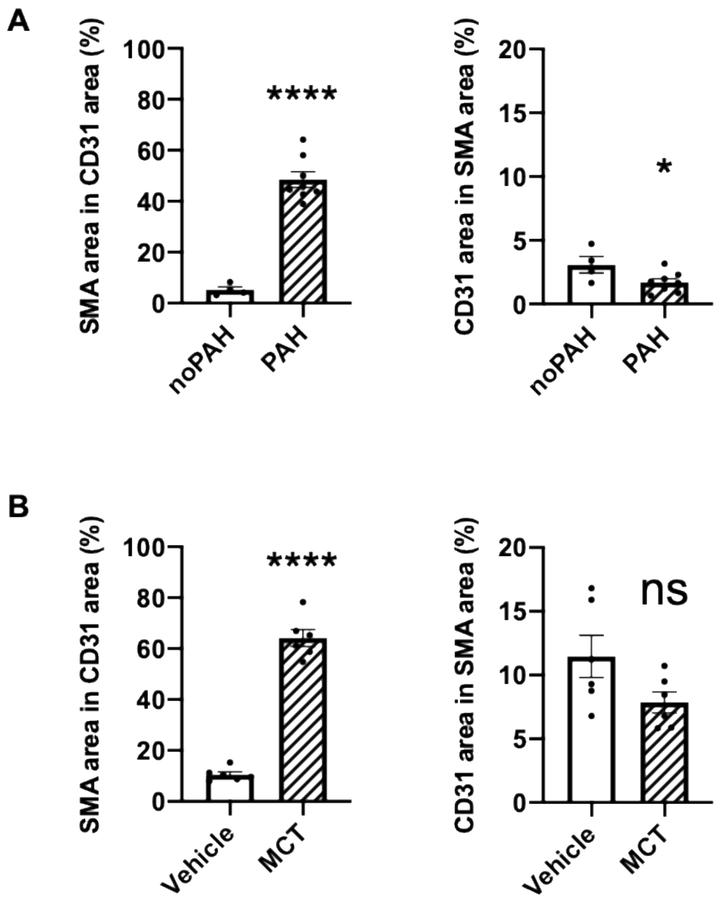


**Supplement Figure S1. Overlap between SMA-positive and CD31-positive cells in the vessels of control and PH vessels.**

A. Colocalization between CD31 and SMA immunofluorescence in the normal and PAH vessels from human patients (n = 4, control; n = 8, PAH. * - $p < 0.05$, **** - $p < 0.0001$).

B. Colocalization between CD31 and SMA immunofluorescence in the normal and PH vessels in MCT-treated rats (n = 6. **** - $p < 0.0001$).

A

B

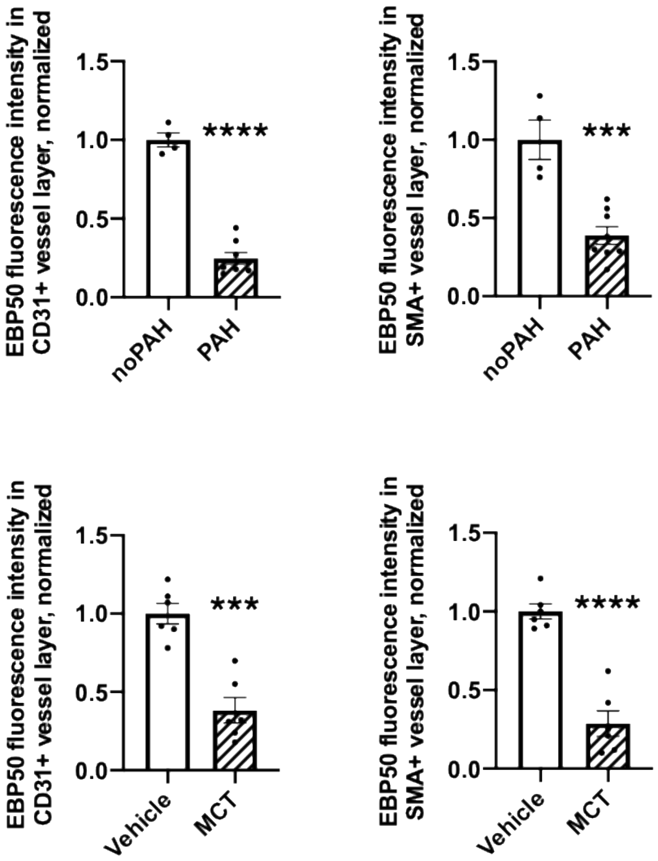


**Supplement Figure S2. EBP50 is downregulated in CD31+ and SMA+ vessel layers in PAH humans and PH rats**

A. Immunofluorescent staining against EBP50 is decreased in intimal (left) and medial (right) layers of PAH lung tissue vessels compared to non-PAH controls. Quantification of the immunofluorescent signal of EBP50 in CD31- or SMA-positive areas per vessel demonstrates a decrease in EBP50 signal in PAH lungs compared controls (n = 4, control; n = 8, PAH. *** - p < 0.001, **** - p < 0.0001).

B. Immunofluorescent staining against EBP50 is decreased in intimal (left) and medial (right) layers of PH lung tissue vessels of monocrotaline-treated rats (MCT) compared to Vehicle control group. Quantification of the immunofluorescent signal of EBP50 in CD31- or SMA-positive areas per vessel demonstrates a decrease in EBP50 signal in PH lungs compared controls (n = 6. *** - p < 0.001, **** - p < 0.0001).

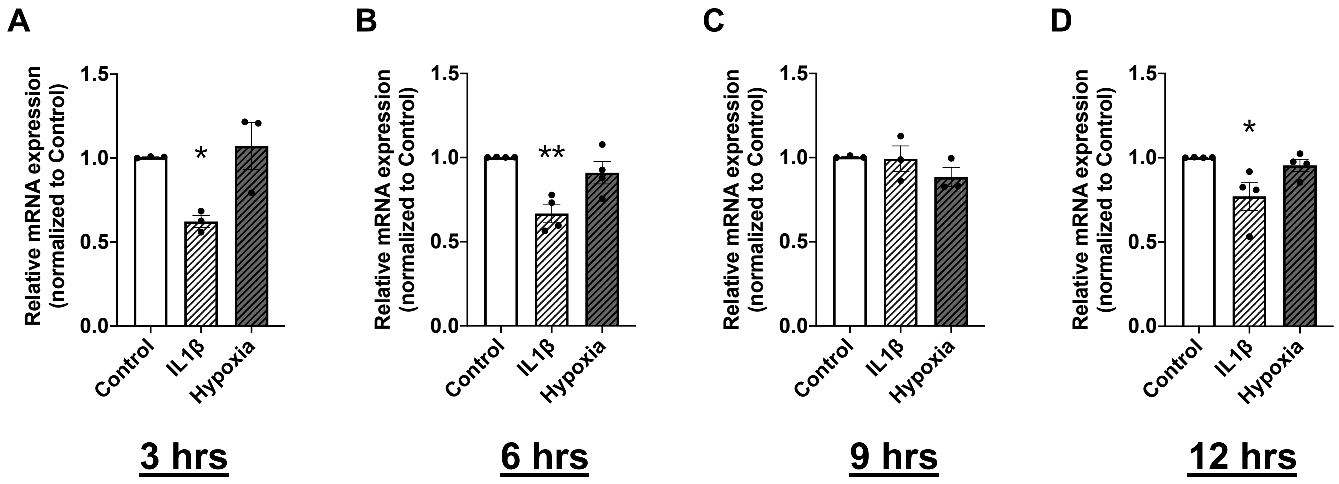


**Supplement Figure S3. EBP50 is downregulated by IL-1β but not hypoxia treatment in HPAECs at early time-points.**

A. EBP50 mRNA level is downregulated following 3 hrs of 10 ng/ml IL-1β treatment and not affected by hypoxia (* - $p < 0.05$, n = 3).
B. EBP50 mRNA level is downregulated following 6 hrs of 10 ng/ml IL-1β treatment and not affected by hypoxia (** - $p < 0.01$, n = 3).
C. EBP50 mRNA level is and not affected by 9 hrs of 10 ng/ml IL-1β treatment or hypoxia (n = 3).
D. EBP50 mRNA level is downregulated following 12 hrs of 10 ng/ml IL-1β treatment and not affected by hypoxia (* - $p < 0.05$, n = 3).

Relative mRNA expression was normalized to β-actin and quantified using ΔΔCt rt-PCR method.

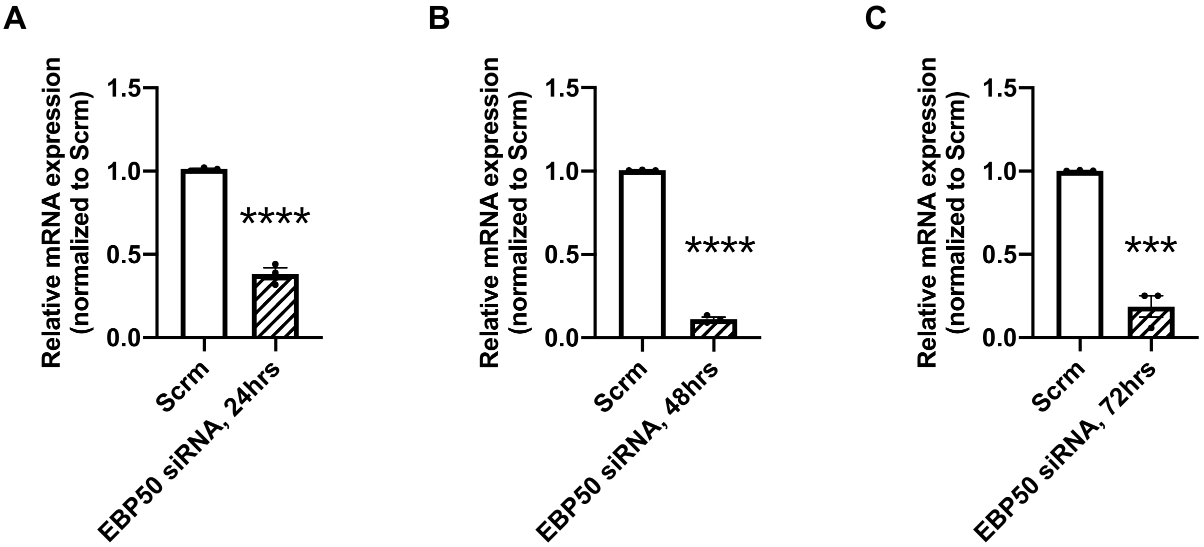


**Supplement Figure S4. EBP50 knockdown efficiency in HPAECs at 24, 48 and 72 hrs post-transfection.**

A. EBP50 mRNA level is decreased by ~ 60% 24 hrs post-transfection with EBP50 siRNA (**** - $p < 0.0001$ from Scrambled, n = 3).
B. EBP50 mRNA level is decreased by ~ 90% 48 hrs post-transfection with EBP50 siRNA (**** - $p < 0.0001$ from Scrambled, n = 3).
C. EBP50 mRNA level is decreased by ~ 80% 72 hrs post-transfection with EBP50 siRNA (*** - $p < 0.001$ from Scrambled, n = 3).

Relative mRNA expression was normalized to 18s and quantified using $\Delta\Delta$Ct rt-PCR method.

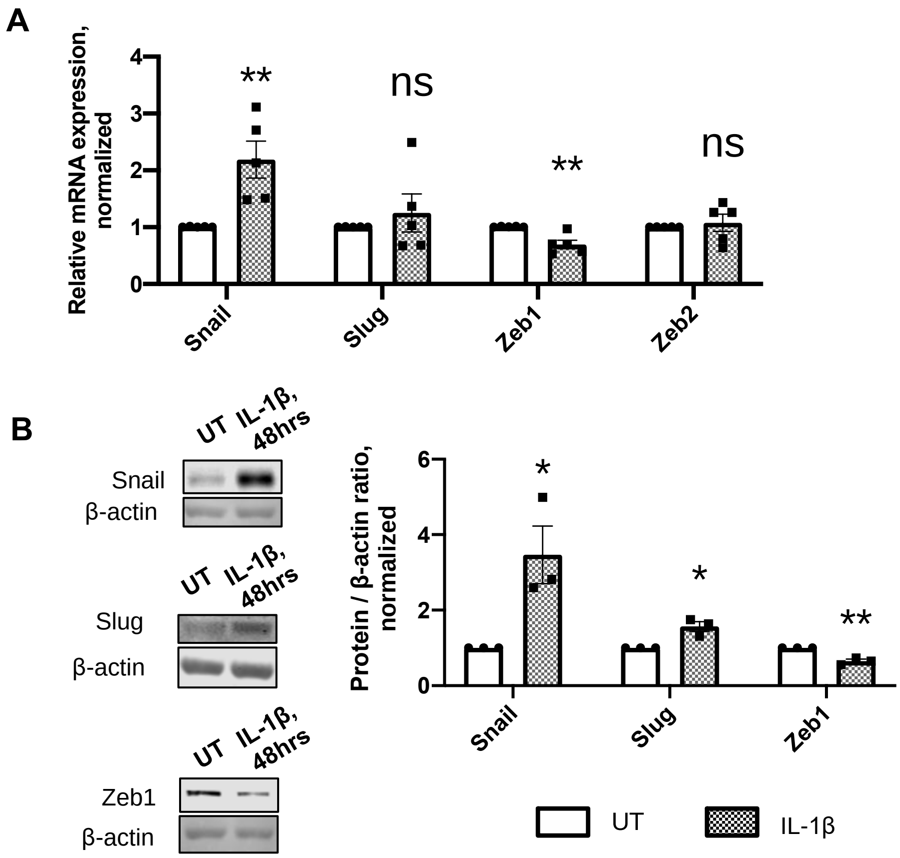


**Supplement Figure S5. IL-1β alters expression of EndMT transcription factors Snail, Slug and Zeb1.**

A. mRNA level of EndMT transcription factor Snail is upregulated following 48 hrs of 10 ng/ml IL-1β treatment, while Zeb1 mRNA is downregulated (** - p < 0.01 from Scrambled siRNA, n = 5).
B. Protein expression of EndMT transcription factors Snail and Slug is upregulated following 48 hrs of 10 ng/ml IL-1β treatment, while Zeb1 is downregulated (* - p < 0.05, ** - p < 0.01 from Scrambled siRNA, n = 3).

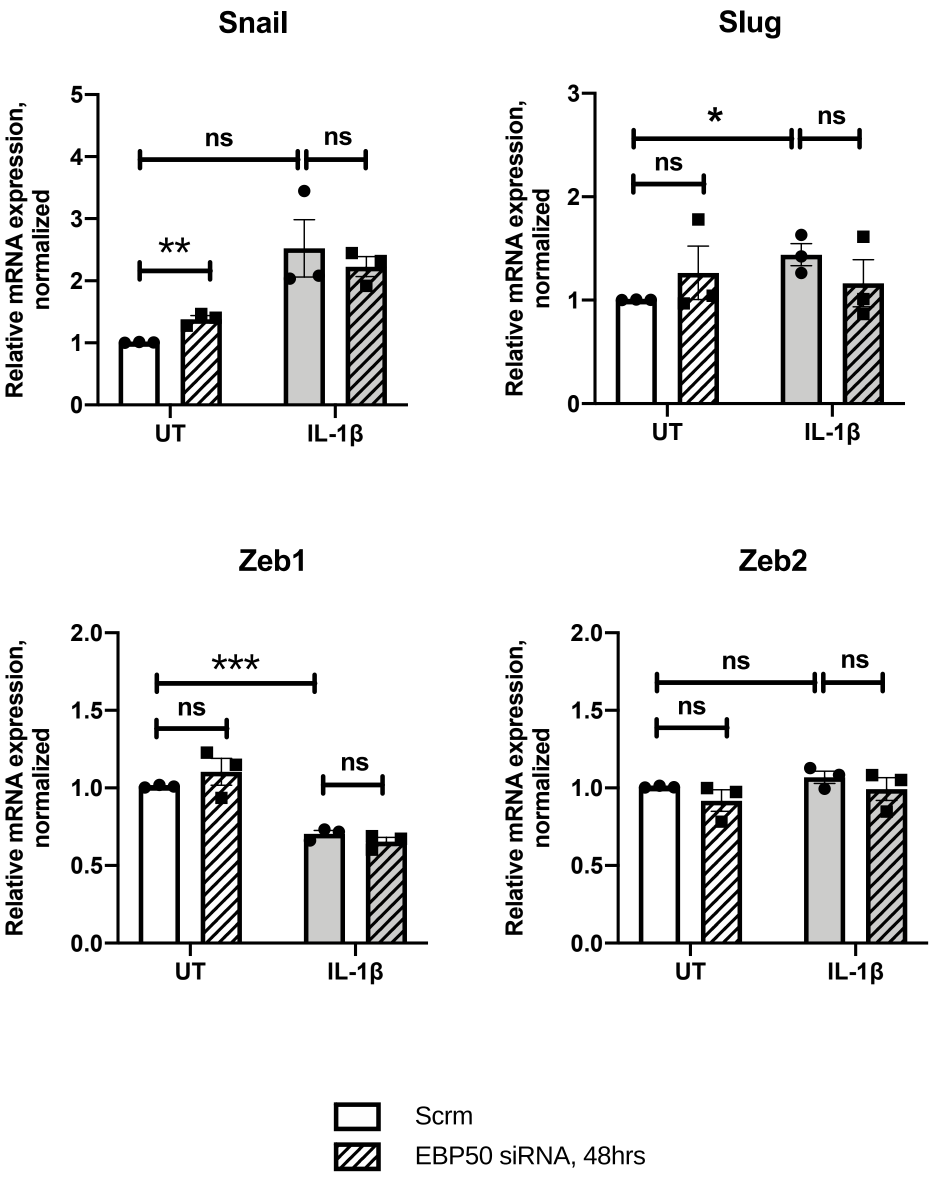


**Supplement Figure S6. EBP50 knockdown has no effect on IL-1β-induced changes in expression of EndMT transcription factors.**

A. Snail expression is upregulated by EBP50 knockdown and unaffected by IL-1β.
B. Slug expression is upregulated by IL-1β and unaffected by EBP50 knockdown.
C. Zeb1 expression is downregulated by IL-1β and unaffected by EBP50 knockdown.
D. Zeb2 expression is unaffected by EBP50 knockdown or IL-1β.

* - $p < 0.05$, *** - $p < 0.001$, n = 3. mRNA normalized to 18s was quantified using ΔΔCt rt-PCR method.

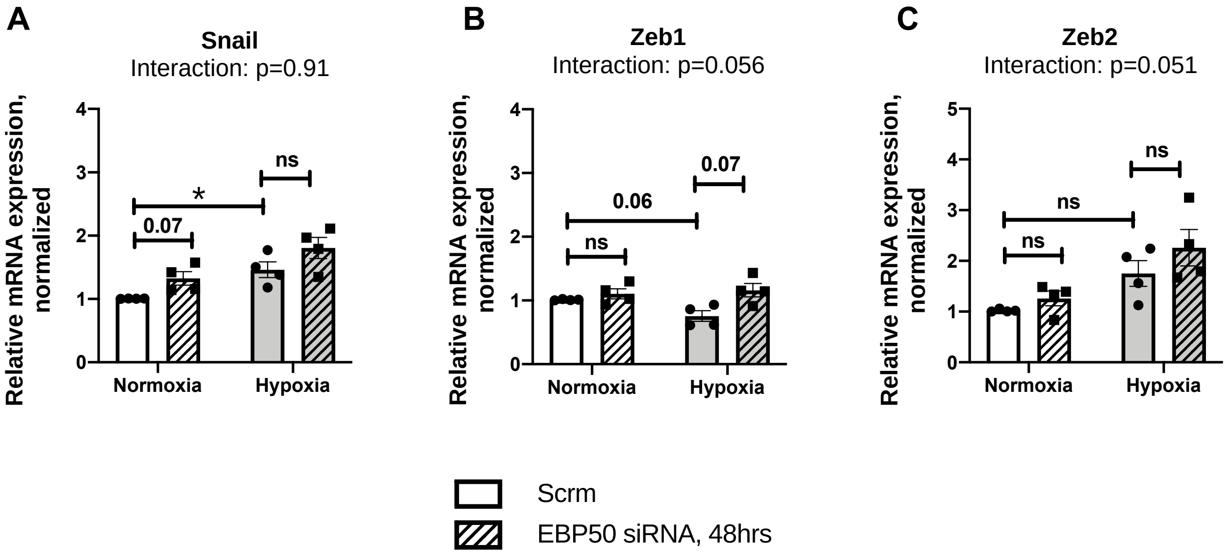


**Supplement Figure S7. EBP50 knockdown does not affect hypoxia-induced changes in EndMT transcription factors Snail, Zeb1, or Zeb2 expression.**
Expression of Snail (A), Zeb1 (B) and Zeb2 (C) was not differentially affected by a combination of hypoxia and EBP50 knockdown at 48 hrs.
N = 4, * - $p < 0.05$.
mRNA normalized to 18s was quantified using $\Delta\Delta$Ct rt-PCR method.

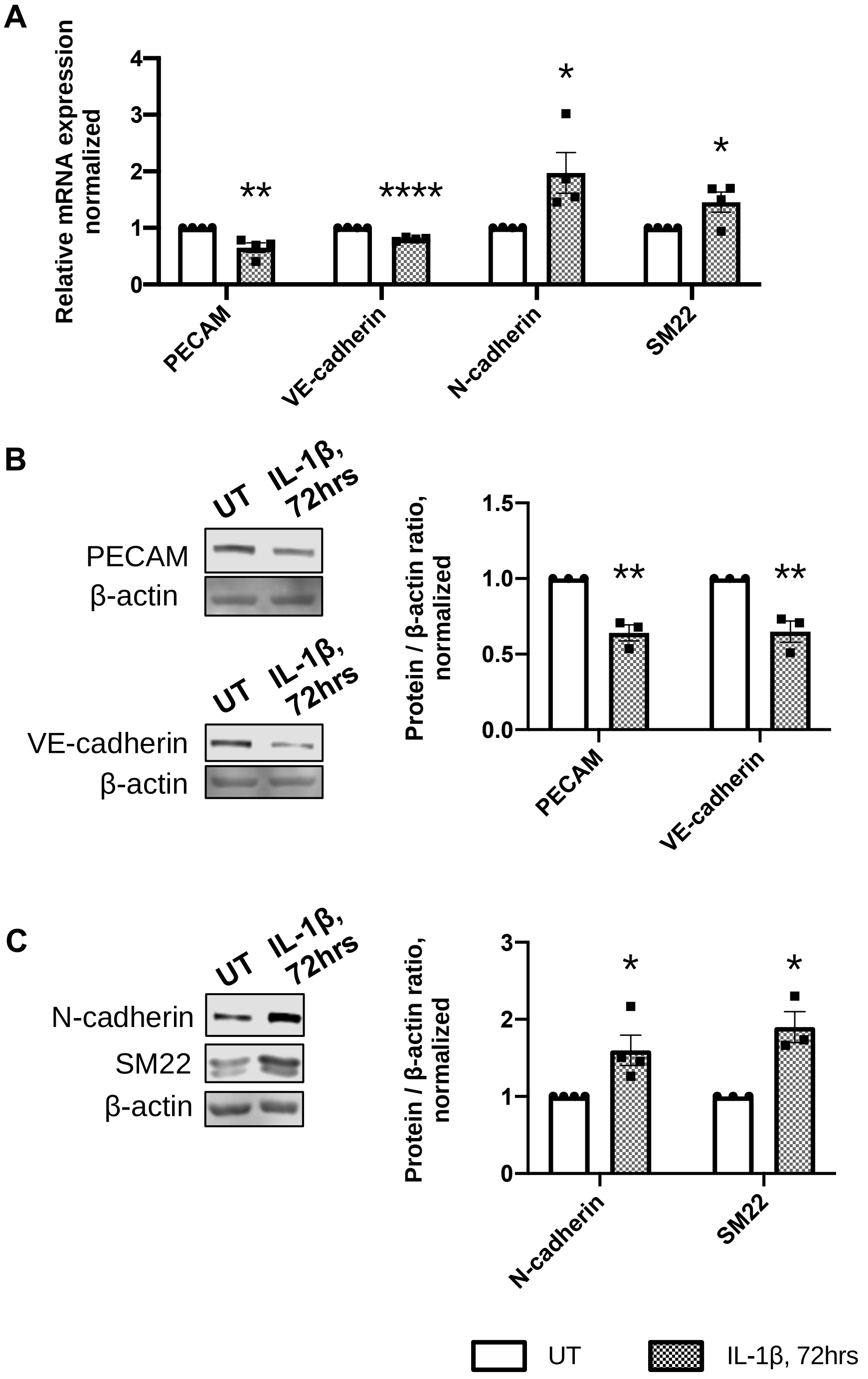


**Supplement Figure S8. IL-1β promotes a loss of endothelial markers and a gain of mesenchymal markers by the pulmonary endothelial cells.**

A. mRNA expression of endothelial markers PECAM and VE-cadherin is downregulated, while mesenchymal markers N-cadherin and SM22 are upregulated following 72 hrs of 10 ng/ml IL-1β treatment (* - $p < 0.05$, ** - $p < 0.01$, **** - $p < 0.0001$ from Scrambled siRNA, n = ). mRNA normalized to 18s was quantified using ΔΔCt rt-PCR method.
B. Representative Western immunoblots (left) and densitometric quantification (right) of PECAM and VE-cadherin demonstrate a decrease in protein expression following 72 hrs of 10 ng/ml IL-1β treatment (** - $p<0.01$ from Scrambled siRNA, n = 3). Relative protein expression was calculated as a ratio to β-actin level in a respective sample.
C. Representative Western immunoblots (left) and densitometric quantification (right) of N-cadherin and SM22 demonstrate an increase in protein expression following 72 hrs of 10 ng/ml IL-1β treatment (* - $p<0.05$ from Scrambled siRNA, n = 3). Relative protein expression was calculated as a ratio to β-actin level in a respective sample.

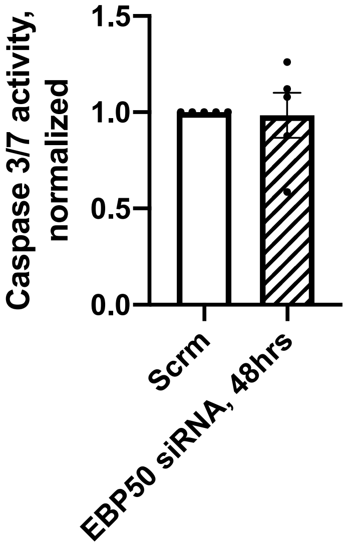


**Supplement Figure S9. 48hrs of EBP50 knockdown does not induce caspase 3/7 cleavage, indicative of apoptosis.**
Luminescent assay of caspase-3/7 activity demonstrated no difference between Scrambled and EBP50 siRNA-transfected HPAECs after 48 hrs of transfection. N=5.

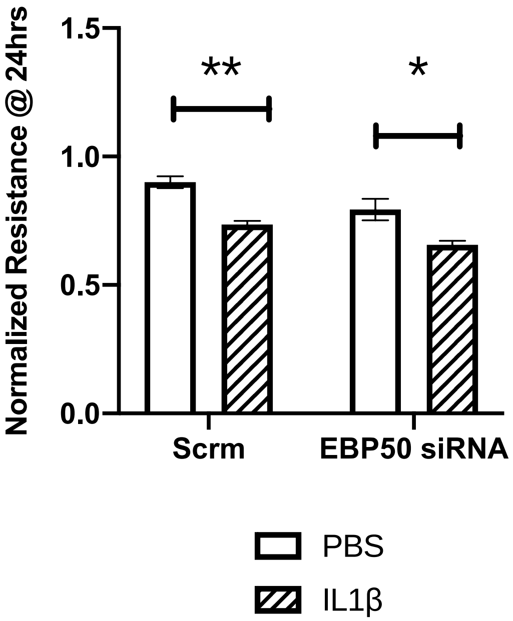


**Supplement Figure S10. EBP50 knockdown did not exacerbate a reduction of endothelial monolayer resistance induced by IL-1β.**
As measured by the ECIS technology, normalized resistance of the HPAECs monolayer was decreased following 24 hrs of treatment with 10 ng/ml IL-1β to a similar extent in Scrambled and EBP50 siRNA-transfected wells.
* - $p < 0.05$, ** - $p < 0.01$, $n = 3$.

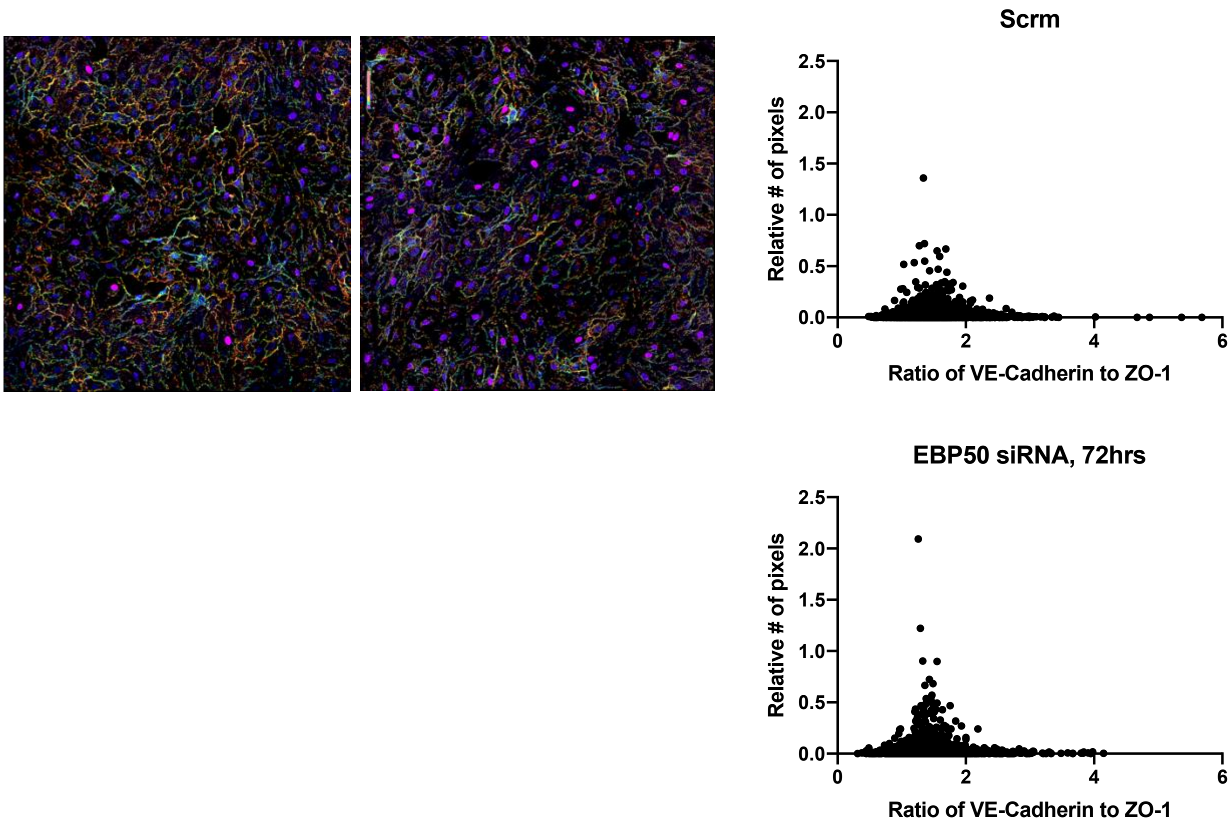


**Supplement Figure S11. EBP50 knockdown did not alter the distribution or abundance of adherens and tight junction proteins VE-cadherin and ZO-1.**
As demonstrated by a representative immunofluorescence image (Panel A) and quantified on the graph on Panel B, EBP50 knockdown did not affect cellular distribution of ZO-1 and VE-cadherin (absence of shift of the ratio curve to the left or right) and did not change the abundance of one protein relative to the other (absence of change in the ratio itself). N=3.

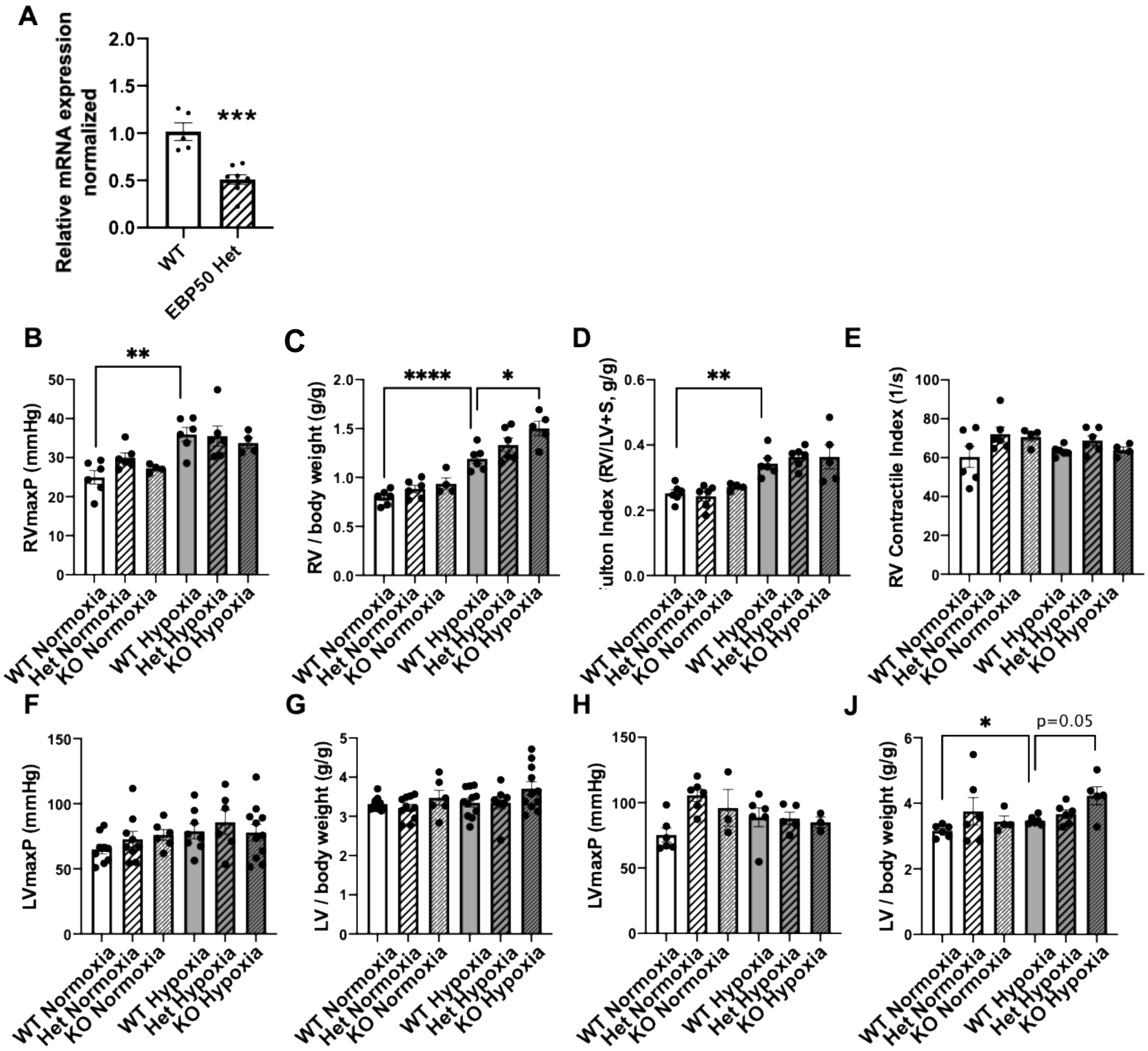


**Supplement Figure S12.**

A. Total lung EBP50 expression in EBP50 Heterozygous mice is decreased by ~ 50% compared to wild-type (*** - $p < 0.001$ from Scrambled, n = 5-8). Relative mRNA expression was normalized to 18s and quantified using ΔΔCt rt-PCR method.

B-J: For a period of chronic hypoxia (4 weeks) treatment, experimental animals were placed in an environment with $FiO_2$ = 10%. At the end of the treatment period, pressure-volume loop measurements were acquired through high-fidelity admittance catheter and hemodynamic parameters were calculated.

B. EBP50 Het Hypoxia male mice did not exhibit an exacerbated RV maximum pressure compared to WT Hypoxia group; n = 4-6.

C. EBP50 Het Hypoxia male mice did not exhibit an exacerbated RV hypertrophy calculated as a ratio of RV weight (g) to body weight (g) compared to WT Hypoxia group; n = 4-6.

D. EBP50 Het Hypoxia male mice did not exhibit an exacerbated RV hypertrophy (increased Fulton Index) compared to WT Hypoxia group; n = 4-6.

E. EBP50 Het Hypoxia male mice did not exhibit an exacerbated RV Contractile Index compared to WT Hypoxia group; n = 4-6.

F. EBP50 Het Hypoxia female mice did not exhibit an exacerbated LV maximum pressure compared to WT Hypoxia group; n=6-11.

G. EBP50 Het Hypoxia female mice did not exhibit an exacerbated LV hypertrophy calculated as a ratio of LV weight (g) to body weight (g) compared to WT Hypoxia group; n=6-11.

H. EBP50 Het Hypoxia male mice did not exhibit an exacerbated LV maximum pressure compared to WT Hypoxia group; n = 3-6.

I. EBP50 Het Hypoxia male mice did not exhibit an exacerbated LV hypertrophy calculated as a ratio of LV weight (g) to body weight (g) compared to WT Hypoxia group; n = 4-6.

* - $p < 0.05$, ** - $p < 0.01$, **** - $p < 0.0001$.
