## Supplemental Methods for "The Role of EBP50 in Regulating Endothelial-To-Mesenchymal Transition in Pulmonary Hypertension"

Anastasia Gorelova et al.

### Supplemental Material

Human tissue samples were obtained through the University of Pittsburgh Tissue Processing Core, Boston Children's Hospital, Brigham and Women's Hospital, and University of California, Los Angeles, and approved by the University of Pittsburgh, Boston Children's Hospital, Brigham and Women's Hospital, and University of California, Los Angeles Institutional Review Boards. Human pulmonary artery endothelial cells from de-identified PAH patients and non-diseased subjects were obtained from the University of Pittsburgh VMI Cell Processing Core and Pulmonary Hypertension Breakthrough Initiative (PHBI) in agreement with the University of Pittsburgh and PHBI protocols. All animal experiments were approved by and conducted in accordance with the University of Pittsburgh Institutional Animal Care and Use Committee.

**Human pulmonary endothelial cells isolation:** De-identified early-passage (3-8) human PAECs from non-diseased subjects and patients with PAH were provided by the University of Pittsburgh Cell Processing Core and Pulmonary Hypertension Breakthrough Initiative (see **Supplementary Table 2** for human subject characteristics). De-identified lung tissue specimens for cell isolation by Cell Processing Core were obtained from the University of Pittsburgh Tissue Processing Core in accordance to the protocols approved by the University of Pittsburgh Institutional Review Board (IRB) and the Committee for Oversight of Research and Clinical Training Involving Decedents (CORID). Cell isolation and maintenance were performed as described previously with modifications for endothelial cells [1]. Briefly, first and second order pulmonary arteries were extensively cleaned from fat and adventitia and perfused with 0.5% Trypsin (Gibco, Carlsbad, CA) and 2.5 mg/ml Collagenase I (Worthington Biochemical Corporation, Lakewood, NJ). Resulting endothelial cell-rich perfusate was collected and plated on cell culture-treated plastic dishes. The purity of the resulting culture maintained in VascuLife VEGF Endothelial Medium (Lifeline Cell Technology, Frederick, MD) was confirmed by a combination of vWF immunofluorescent staining (vWF antibody, Abcam, United Kingdom) and visual identification.

**Mouse pulmonary endothelial cells isolation:** Pulmonary endothelial cells were isolated as previously described [2].

**Animals:** All animal experiments were approved by and conducted in accordance with the University of Pittsburgh Institutional Animal Care and Use Committee. To model the monocrotaline (MCT)-induced severe PH, 10-14 weeks old adult male Sprague-Dawley rats (Charles River Laboratories, Hudson, NY) were injected once with 60 mg/kg of monocrotaline and euthanized after 3 weeks [2]. Control animals were maintained under normoxia and injected with vehicle. To model chronic hypoxia-induced PH, 10-12 weeks old adult male C57BL/6 wild type mice (Taconic Biosciences, Germantown, NY and Jackson Laboratory, Bar Harbor, ME) were subjected to continuous normobaric hypoxia (10% $O_2$, BioSpherix, Parish, NY) for three to four weeks (21 – 28 days). Control animals were maintained under normoxia. At the termination of all animal studies, lung samples were snap frozen in liquid nitrogen and stored for later analysis.

**EBP50 Heterozygous and knockout mice:** 9-16 weeks old adult male and female EBP50 Heterozygous mice ($EBP50^{+/-}$, EBP50 Het), EBP50 Knockout mice ($EBP50^{-/-}$, EBP50 KO), or wildtype (WT) controls bred on C57BL/6 background were subjected to continuous normobaric hypoxia (10% $O_2$, BioSpherix, Parish, NY) for four weeks. At the termination of the study, hemodynamic data was obtained using the 1.2F pressure-volume catheters (Transonic Systems Inc., Ithaca, NY) inserted directly through the right then left ventricle. After the hemodynamic data was obtained, heart and lung samples were collected and snap frozen in liquid nitrogen and stored for later analysis. Hearts were separated into RV and left ventricle (LV) + septum, and Fulton index was calculated as an RV / (LV + septum) ratio.

**Cell culture:** Human pulmonary artery endothelial cells (HPAECs) were obtained from PromoCell GmbH, Germany [Catalog #: C-12241, Lot # 432Z010.2 (donor info: female, causasian 50 y.o.). HPAECs were grown in the Endothelial Cell Growth Medium 2 (EGM-2 medium, PromoCell, Germany) in the absence of heparin [Basal medium, C-22211; Growth Medium 2 Supplement Pack, C-39211] according to manufacturer protocol.

**Cell treatments:** Recombinant human IL-1β was obtained from Peprotech, Inc. (Cat #: 200-01B); lyophilized powder was reconstituted in mQ $H_2O$ with an addition of 0.1% BSA and stored in 100 μl aliquots in the -80 $^oC$ for no longer than one year since the date of preparation. Immediately prior to cell treatment, IL-1β was diluted in cell culture medium to be used at a dose of 10 ng/ml. For hypoxia exposure, cells were cultured in temperature-humidity controlled chamber under $pO_2 = 1\%$, $pCO_2 = 5\%$, with $N_2$ balance at 37$^o$C for 48 – 72 hrs.

**EBP50 knockdown:** To induce EBP50 gene knockdown, HPAECs were transiently transfected with Silencer Select EBP50 siRNA (Cat #: 4390824, ID: s17920, Life Technologies, Carlsbad, CA) using the Lipofectamine 3000 transfection reagent (Cat #: L3000015, Invitrogen, Carlsbad, CA) according to the manufacturer's protocol for 24 – 72 hrs. Silencer Select Negative Control No. 1 siRNA (Cat #: 4390843, Invitrogen, Carlsbad, CA) was used as a control for potential off-target effects of the transfection procedure.

**Quantitative real-time PCR:** Levels of mRNA transcripts were quantified as previously described [3]. Briefly, total RNAs were extracted from rat and mouse lung tissues, isolated pulmonary endothelial cells, and from HPAECs using the RNeasy Plus Mini Kit (Qiagen, Germany) according to the manufacturer's protocol. Total RNA (0.5 μg) was reverse transcribed to cDNA using Superscript III reverse transcriptase (Life Technologies, Carlsbad, CA) following the manufacturer's protocol. TaqMan probe and Universal PCR Master Mix and SLC9A3R1, SNAI1, SNAI2, Zeb1, Zeb2, CTNNB, PECAM1, CDH5, CDH2, TAGLN, S100A4, FN1, ACTB or 18S primers (Life Technologies, Carlsbad, CA) were used for qPCR in a QuantStudio 6 Flex Real-Time PCR System (Applied Biosystems, Foster City, CA) according to the manufacturer's protocol for 40 cycles. Relative quantification to corresponding control was obtained using the threshold cycle (Ct) method with 18S as the housekeeping gene and relative expression calculated as $2^{-\Delta\Delta Ct}$.

**Western blot:** Western blot experiments were performed as described previously [3]. Total protein (7.5 – 20 μg) from tissue homogenates or cell lysates were added to Tris-glycine SDS sample buffer, boiled, resolved with SDS / PAGE, and transferred onto Trans-Blot nitrocellulose membranes (Bio–Rad, Hercules, CA). Membranes were blocked with the Odyssey Blocking Buffer (LI-COR Biosciences, Lincoln, NE) and incubated with rabbit anti-EBP50 (1:1000 dilution, Invitrogen, Carlsbad, CA), rabbit anti-Zeb1 (1:1000 dilution, Danvers, MA), mouse anti-PECAM (1:1000 dilution, Cell Signaling Technology, Danvers, MA), rabbit anti-VE-cadherin (1:1000 dilution, Cell Signaling Technology, Danvers, MA), rabbit anti-N-cadherin (1:1000 dilution, Cell Signaling Technology, Danvers, MA), rabbit anti-SM22 (1:1000 dilution, Abcam, United Kingdom), mouse anti-vinculin (1:1000, Santa Cruz Biotechnology, Dallas, TX), or mouse anti-β-actin (1:1000, Santa Cruz Biotechnology, Dallas, TX). Membranes were probed with fluorescence-tagged (680 or 800 nm) anti-rabbit or anti-mouse secondary antibodies (1:10,000 dilution, LI-COR Biosciences, Lincoln, NE). Digital membrane scans were obtained using the Odyssey Infrared Imaging system (LI-COR Biosciences, Lincoln, NE). Optical density (OD) of protein-of-interest bands were quantitated using ImageJ software (NIH, USA) and normalized to vinculin or β-actin.

**Electrical Cell-substrate Impedance Sensing:** Barrier function of HPAECs monolayer were quantified with electrical cell-substrate impedance sensing (ECIS; Applied BioPhysics, Troy, NY). Cells were seeded at 3.5 x $10^4$ cells / 0.8 $cm^2$ well, each containing two sets of 20 circular 250 μm diameter active electrodes (8W10E+). Impedance, resistance, and capacitance of the endothelial monolayer was measured continuously over a three-hour period to establish the baseline read, and over 24 hrs to determine effects of IL-1β on endothelial permeability in Scrambled- or EBP50 siRNA-transfected HPAECs.

**EdU incorporation:** To measure the extent of de novo DNA synthesis, Click-iT EdU Pacific Blue Flow Cytometry Assay Kit (Cat #: C10418, Invitrogen, Carlsbad, CA) was used according to the manufacturer's protocol.

**Statistics:** All results are reported as mean ± SEM. For in vitro studies, where normality was not tested, normal distribution was assumed. Unpaired two-tailed Student's t-test was used for comparison between two groups. To test the simultaneous effect of EBP50 and hypoxia in vitro and in vivo, we used a test of interaction from a two-way ANOVA with bootstrapping. All variables tested, with the exception of Fulton index, had passed the test for normality. Distribution of Fulton Index values departed from normality, however, the natural log transformation did not impact the analysis (p value of interaction after log transformation = 0.15). For all analyses, $p < 0.05$ was deemed statistically significant. GraphPad Prism Software v7 and Stata 16.1 were used for data analyses.


1. Goncharov, D. A. *et al.* Mammalian target of rapamycin complex 2 (mTORC2) coordinates pulmonary artery smooth muscle cell metabolism, proliferation, and survival in pulmonary arterial hypertension. *Circulation* **129**, 864–74 (2014).
2. Bertero, T. *et al.* Systems-level regulation of MicroRNA networks by miR-130/301 promotes pulmonary hypertension. *J. Clin. Invest.* **124**, 3514–3528 (2014).
3. Ghouleh, I. Al *et al.* Endothelial Nox1 oxidase assembly in human pulmonary arterial hypertension; driver of Gremlin1-mediated proliferation. *Clin. Sci. (Lond).* **131**, 2019–2035 (2017).
